# A genome-wide atlas of meiotic recombination intermediates reveals distinct modes of DNA repair that direct crossovers away from transcriptionally marked genes

**DOI:** 10.64898/2026.03.26.714455

**Authors:** Callum Henfrey, Emily Print, Gang Zhang, Robert Hinch, Isabella Maudlin, Daniela Moralli, Benjamin Davies, Peter Donnelly, Anjali Gupta Hinch

## Abstract

Crossovers are essential for accurate chromosome segregation in meiosis. Yet the programmed DNA double-strand breaks that initiate them frequently occur in genes and pose a risk to transcription required for gametogenesis. How meiotic cells reconcile these competing demands has remained unclear. Here, we generate a genome-wide *in vivo* atlas of meiotic recombination intermediates across ∼42,000 hotspots by mapping repair proteins BLM, HFM1, and RPA in wild-type and genome-engineered mutant mouse testes. These maps reveal two distinct modes of break repair: a fast-resolving class with short-lived intermediates that are repaired predominantly as non-crossovers, and a slower class with persistent intermediates that give rise to nearly all crossovers. Fast-resolving hotspots occur almost exclusively within a deeply conserved set of ∼4,500 genes marked by structural and chromatin features established during an early stage of meiotic transcription. This transcriptional memory predicts repair fate with high accuracy across mouse subspecies and sexes. Across widely diverged mammals, including humans and cattle, orthologous genes show similar crossover suppression. Our findings reveal an early bifurcation between crossover and non-crossover repair that is governed by the transcriptional context of meiotic breaks. Together, they establish an evolutionarily conserved principle in which crossovers are directed away from transcriptionally important genes, thereby safeguarding gene function and shaping their evolution.

## Main Text

Meiotic recombination is essential for fertility and for generating genetic diversity in sexually-reproducing species^1,2^. By forming crossovers between homologous chromosomes, it ensures their accurate segregation and reshuffles genetic variants that drive evolutionary change^1,3^. Recombination is initiated by the programmed formation of hundreds of DNA double-strand breaks (DSBs), which in humans and other species cluster within narrow genomic regions known as “hotspots”^4^. Only a small fraction of these breaks – approximately 10% – are resolved as crossovers^5^. The majority are resolved as non-crossovers through homologous recombination without reciprocal exchange^5^.

Repairing DSBs in a genome that must simultaneously orchestrate extensive transcription during meiosis presents a fundamental challenge. DSB repair is mutagenic, disrupts ongoing transcription, and risks forming non-canonical DNA:RNA structures or transcription-repair collisions^6-9^. Yet in humans and mice, meiotic DSBs are enriched within gene bodies^10,11^; regions where interference with transcription may be particularly detrimental.

Crossovers impose additional constraints: their multi-protein intermediates tether homologous chromosomes for prolonged periods^1^, potentially restricting transcriptional access at associated loci. Notwithstanding the increased rate of DSBs in genes, population-based recombination maps have reported reduced crossover frequency in genic regions and chromatin states associated with transcription in testes, although these broad annotations do not identify the relevant developmental stage nor distinguish genes from wider regulatory domains^12-15^. Moreover, because transcription in testes is extensive, with over 80% of genes expressed^16-19^, such correlations provide limited insight into regulation of meiotic DNA break repair.

A central unresolved question is therefore how meiotic cells control the timing and outcomes of break repair across the genome. Addressing this question requires a comprehensive and finely detailed understanding of recombination intermediates that capture the transient states underlying repair-pathway choice. The Bloom syndrome helicase BLM acts on DNA repair intermediates to regulate whether they are resolved with or without crossovers^20,21^. We have generated the first genome-wide *in vivo* maps of BLM, together with additional recombination factors HFM1 and RPA, across ∼42,000 hotspots in wild-type and genetically-engineered mutant mouse testes. These maps reveal a previously unrecognised regulatory architecture in which transcriptional history shapes the kinetic processing and outcome of meiotic breaks. In doing so, they uncover unexpected genome-wide coordination between meiotic recombination and transcription.

### Two temporally distinct modes of meiotic break repair revealed by BLM

Meiotic recombination proceeds through a defined sequence of molecular events (Figure 1A). In most mammals, DSBs are positioned by the histone methyltransferase PRDM9 through its binding of specific DNA sequence motifs^22-24^. A few hundred of these sites are broken by SPO11^25^ in a typical meiosis. Resection generates single-stranded DNA (ssDNA) overhangs bound by RPA^26-28^, which interacts with BLM^29^. The recombinase DMC1, supported by RAD51, mediates strand invasion into the homologous chromosome, generating nascent strand-exchange intermediates (D-loops), some of which are stabilised by HFM1 (Mer3)^30-32^. BLM acts on strand-exchange intermediates to regulate their ultimate resolution^1,33^. Whilst some correlates of crossover formation have been identified^34-38^, the regulatory logic governing how long strand-exchange intermediates persist, and how they are ultimately resolved, remains poorly understood.

**Figure 1.**
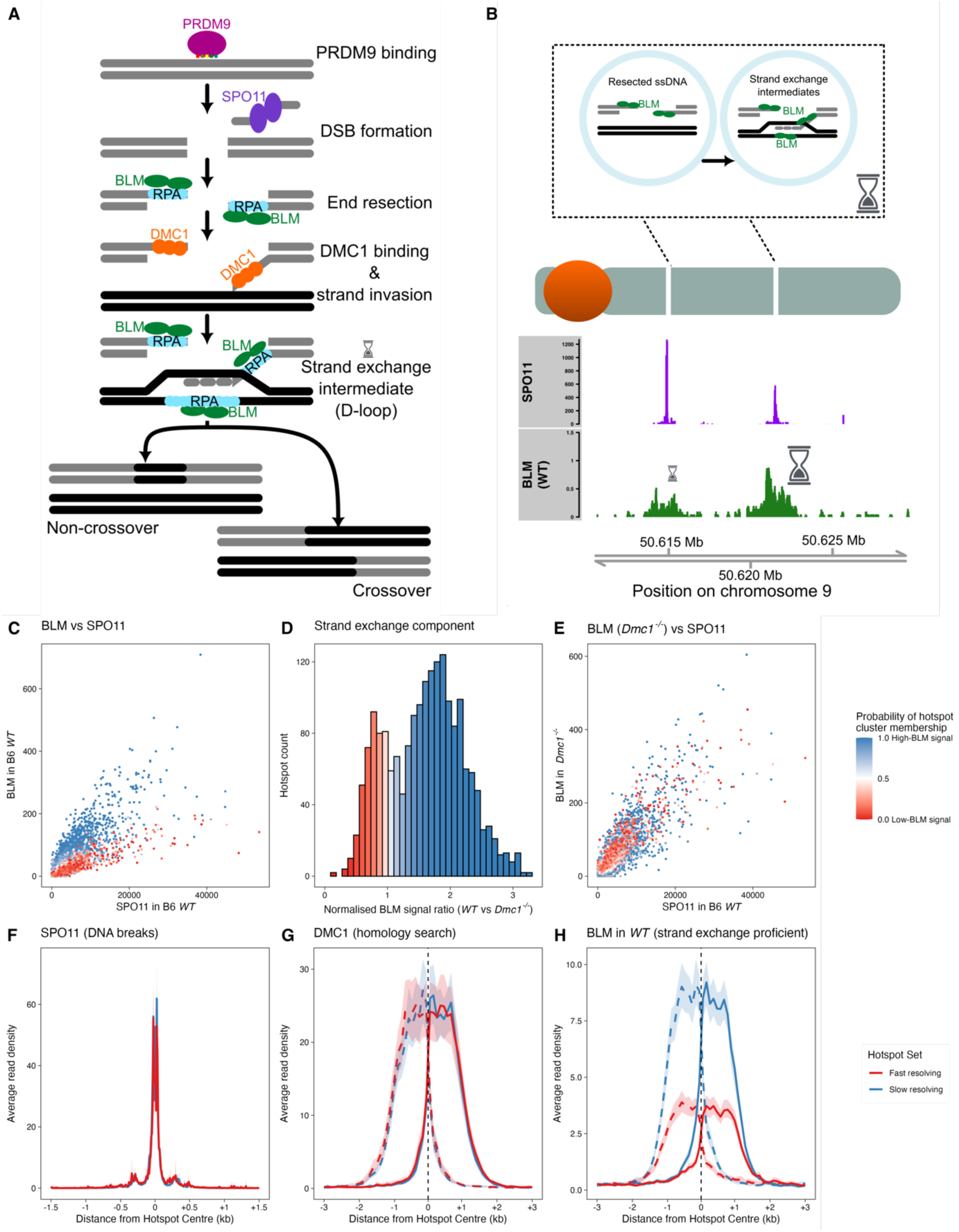
Genome-wide maps of BLM reveal two temporally distinct modes of meiotic DNA repair. **(A)** SPO11 induces programmed DNA double-strand breaks (DSBs), which are resected to generate single-stranded DNA (ssDNA) coated by RPA, which interacts with BLM. DMC1 replaces RPA to mediate homology search and strand invasion, forming strand-exchange intermediates (D-loops). BLM binds these intermediates, which are ultimately resolved as crossovers (∼10%) or non-crossovers (∼90%). **(B)** (Top) BLM ChIP-SSDS signal reflects BLM association with resected ssDNA and strand-exchange intermediates and is impacted by their lifespans. (Bottom) Representative hotspots showing SPO11 oligos (measuring DSBs, purple) and BLM ChIP-SSDS signal (read counts per million) in WT mice, illustrating variability in BLM signal relative to break frequency. **(C)** Scatterplot of BLM signal versus SPO11-oligo counts in WT mice across 16,658 autosomal hotspots (each point represents one hotspot). Points are coloured according to inferred class assignment (high, blue; low, red). Hotspots separate into two distinct classes with systematically higher and lower BLM signal relative to break frequency. **(D)** Histogram of the strand-exchange–dependent component of BLM signal, inferred through comparison between WT and *Dmc1*^-/-^ mice that lack strand exchange (Supplementary Text). The distribution reveals two classes of hotspots with a high (blue) and low (red) strand-exchange component. Hotspots with sufficient signal to infer class membership (>50 ChIP-SSDS reads in *Dmc1*^-/-^, n=1,902) included; bins coloured by the mean inferred class probability of hotspots in each bin. **(E)** Scatterplot as in (C) for BLM signal in *Dmc1^-/-^* mice versus WT SPO11-oligo counts. This demonstrates that the strand-exchange component of BLM signal disappears in *Dmc1^-/-^*, which eliminates the difference between hotspot classes. **(F)** Average SPO11-oligo density in matched sets of high (blue) and low (red) BLM signal hotspots (10 bp smoothing), showing similar DSB levels between classes. **(G)** Average DMC1 signal in matched hotspots in WT mice (100 bp smoothing), shown for the top (dashed) and bottom (solid) strands. They demonstrate comparable strand-invasion efficiency between the two classes. **(H)** Average BLM signal in matched hotspots in WT mice (100 bp smoothing), shown for the top (dashed) and bottom (solid) strands. The ∼2.3-fold difference between classes reflects differential persistence of strand-exchange intermediates.

To examine this directly, we mapped BLM bound to ssDNA genome-wide in wild-type mouse testes (Figure 1B, Table S1) using ChIP-seq specialised for single-stranded DNA (ChIP-SSDS)^39^. BLM peaks overlapped known DNA break hotspots (Figure 1B) and BLM signal across ∼18,000 previously identified hotspots was strongly correlated with independent break-associated assays (SPO11, RPA, and DMC1; Figure S1A-C), validating BLM ChIP-SSDS as a sensitive reporter of recombination intermediates.

At each hotspot, BLM signal reflects three factors: the number of breaks formed, the extent of BLM loading onto repair intermediates, and the persistence lifespan of those intermediates. Among hotspots with similar DNA break frequency, we observed substantial variation in BLM signal (Figure 1C). This indicates that differences in BLM signal between hotspots cannot be attributed solely to variation in DNA break frequency and may reflect differences in BLM loading or intermediate lifespan.

Because BLM binds both before and after strand invasion (Figure 1A–B), we asked whether the observed variation reflects differences in strand-exchange intermediates. To test this, we generated an equivalent map in *Dmc1*^-/-^ mice, in which strand-exchange intermediates do not form^40,41^ (Figure S1D-G, Table S1). Because *Dmc1*^-/-^ mice fail to undergo strand invasion, the ratio of BLM signal in wild-type relative to *Dmc1*^-/-^ testes isolates the strand-exchange–dependent component of BLM signal (Supplementary Text, Figure S1D,G).

Strikingly, the distribution of this strand-exchange–dependent component revealed that autosomal hotspots segregate into two sharply defined classes (Figure 1D, Figure S1G): one comprising ∼20% of hotspots with low strand-exchange–dependent BLM signal, and the remainder with ∼2.3-fold higher signal on average. Importantly, this class separation relative to break frequency was evident in the wild-type (Figure 1C) but collapsed in the absence of strand exchange (Figure 1E), establishing that class separation is dependent on successful strand exchange.

Differences in the strand-exchange component could arise from variation in BLM loading or strand invasion efficiency rather than intermediate lifespan. To evaluate these possibilities, we compared these features between hotspot classes. Hotspots matched for break frequency (STAR Methods, Figure 1F) showed similar resection length and extent of BLM loading (Figure 1G-H, Figure S1H,J-L). The time taken to locate the homologous template, assessed via the lifespan of DMC1-bound intermediates^36^, was also similar between the hotspot classes, indicating that strand-exchange intermediates form with comparable efficiency (Figure 1G, Figure S1I). Thus, the distinction between hotspot classes does not arise from differences in BLM loading or strand invasion efficiency.

Instead, orthogonal assays indicate that the distinction arises from differences in the lifespan of strand-exchange intermediates. END-seq, a protein-independent assay that reports DNA structures associated with strand-exchange intermediates^42^, showed the same class separation in wild-type testes. This separation was absent in *Dmc1*^-/-^ and *Hop2*^-/-^ mice, both of which lack strand exchange (Figure SJ-L). RPA ChIP-SSDS showed the same pattern: separation in wild type but not in *Dmc1*^-/-^ testes (Figure S1M–P, Table S2). These independent assays rule out differences in BLM loading or antibody artefacts (Supplementary Text). Instead, they show that the two hotspot classes differ in the lifespan of strand-exchange intermediates in wild-type mice (Supplementary Text).

We have therefore identified two temporally distinct modes of meiotic break repair, defined by short-lived (“fast-resolving”) and longer-lived (“slow-resolving”) strand-exchange intermediates.

### Strand-exchange intermediate lifespan marks an early bifurcation between crossover and non-crossover outcomes

To relate fast- and slow-resolving hotspots to eventual repair outcomes, we generated BLM maps in hybrid B6×CAST testes (Table S1) and estimated its signal in ∼24,000 previously identified hotspots^36^ (Figure S2A-F). Equivalent maps in *Dmc1*^-/-^ hybrids were generated to isolate the post–strand-exchange component, as above (Figure S2A-F). These hybrids carry a wild-type *Prdm9*^CAST^ allele and a genetically-engineered *Prdm9* allele containing a DNA-binding domain found in human populations (hereafter, *Prdm9*^HUM^)^11^. In addition to specifying break sites, PRDM9 binding to the homologous template promotes homologue engagement and is essential for efficient pairing^11^. In hybrids, PRDM9 binding can be symmetric (both homologues bound) or asymmetric (only the broken homologue bound), and asymmetric hotspots frequently fail to engage the homologous chromosome^11^. To avoid confounding effects from variable homologue engagement, all analyses were restricted to symmetric hotspots.

Hotspots in the hybrid genome again segregated robustly into fast- and slow-resolving classes (Figure 2A, Figure S2A-G). To assess their recombination outcomes, we used two complementary datasets: single-sperm DNA sequencing, which identifies individual crossover events at high resolution in male meiosis^36^, and pedigree-based recombination maps, which report crossover and non-crossover transmission across meioses in both sexes^37^.

**Figure 2.**
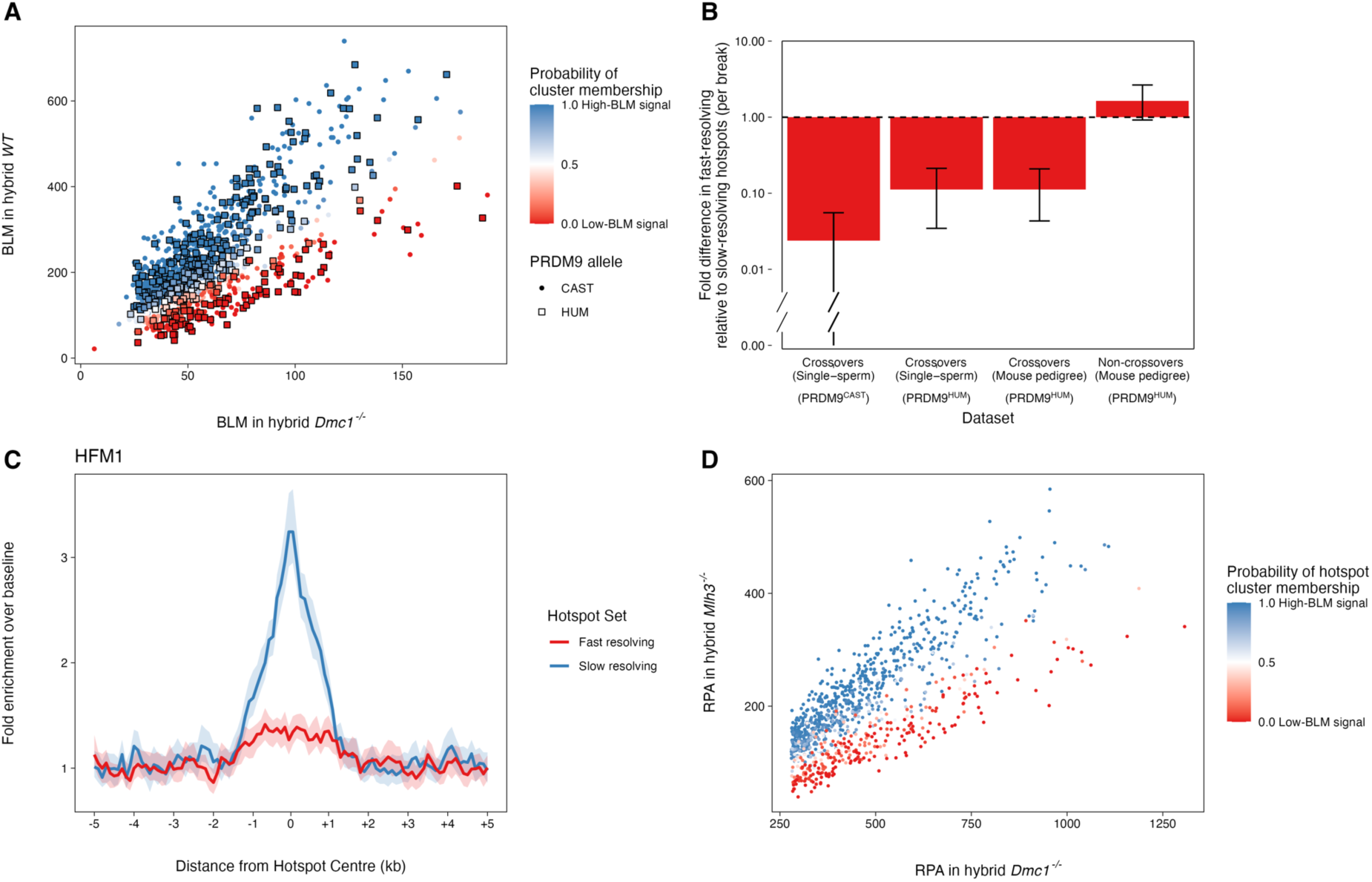
Strand-exchange intermediate persistence marks an early divergence between crossover and non-crossover outcomes. **(A)** Scatterplot of BLM signal in wild-type hybrid relative to the *Dmc1*^-/-^ hybrid for symmetric hotspots. Each point represents one hotspot and is coloured by its inferred class assignment (slow-resolving, blue; fast-resolving, red; n=866). The separation of hotspots reflects differences in strand-exchange intermediate persistence similar to B6. Hotspots controlled by PRDM9^CAST^ (circles) and PRDM9^HUM^ (squares) are indicated. **(B)** Fast-resolving hotspots are strongly depleted for crossovers relative to slow-resolving hotspots, whereas non-crossover outcomes are comparatively enriched. At first glance it may seem paradoxical that sites with higher BLM signal (slow-resolving hotspots) produce more crossovers, given that BLM promotes non-crossover repair^20,21^. However, recall that the higher BLM signal reflects longer persistence of strand-exchange intermediates rather than increased BLM loading. Fold-differences in recombination outcomes normalised for estimated DNA break frequency (using RPA signal in *Dmc1*^-/-^ as proxy^43^) are shown. Crossovers identified by single-sperm sequencing (left), and crossovers and non-crossovers from a *Prdm9*^HUM^ pedigree (right) are shown (data from^36^ and^37^). Non-crossover counts were corrected for local polymorphism density (Figure S2H). Error bars indicate 95% confidence intervals (bootstrap n=2000). **(C)** Average HFM1 ChIP-SSDS signal in matched sets of B6 hotspots from slow-resolving (blue) and fast-resolving (red) classes (matching as per Figure 1), indicating preferential stabilisation of strand-exchange intermediates in slow-resolving hotspots. **(D)** Scatterplot of RPA ChIP-SSDS signal in *Mlh3^-/-^* hybrid versus *Dmc1^-/-^*hybrid testes. Hotspot class separation is preserved in the absence of MLH3-dependent crossover designation, indicating that separation of breaks into fast and slow repair modes precedes crossover specification.

Crossover frequency was markedly lower at fast-resolving than at slow-resolving hotspots (Figure 2B). In sperm, crossovers were suppressed 18-fold across all hotspots, with 42-fold suppression in PRDM9^CAST^ (95% CI=[18.0,∞)) and 9-fold in PRDM9^HUM^ (95% CI=[5,29]) hotspots. The pedigree-based recombination maps^37^ contain sufficient data only for PRDM9^HUM^ hotspots. Crossovers from both sexes showed comparable reduction in these hotspots (9-fold reduction; 95% CI=[5,23]), indicating that crossover suppression at fast-resolving hotspots is a shared feature of both sexes. Together, these results imply that only ∼1% of breaks in fast-resolving hotspots are repaired as crossovers.

In contrast with crossovers, non-crossovers were not suppressed and were potentially enriched (1.6-fold; 95% CI=[0.92, 2.65], Figure 2B) at fast-resolving hotspots, showing that breaks in fast-resolving hotspots are overwhelmingly repaired as non-crossovers. The presence of this behaviour in hotspots marked by PRDM9^HUM^ (Figure 2B), which has not evolved in mouse, demonstrates that differences in strand-exchange intermediate lifespan and outcomes reflect an intrinsic property of the recombination process rather than local evolutionary adaptations.

The predominance of non-crossover repair at fast-resolving hotspots suggests that these events contribute to homology search and early pairing. Two possibilities could explain differences in strand-exchange intermediate turnover^44-46^: One is that intermediates in fast-resolving hotspots are resolved early with minimal stabilisation, potentially before homologous chromosomes are fully synapsed. The other is that intermediates at both fast-and slow-resolving hotspots initially persist, and only later—after all autosomes are synapsed in pachytene—is a subset designated for crossover formation.

To distinguish between these possibilities, we generated genome-wide maps of HFM1 (Table S3), a helicase that protects and stabilises nascent recombination intermediates^30-32^. HFM1 acts early in the ZMM-stabilisation pathway. Unlike BLM, which associates broadly with strand-exchange intermediates, HFM1 marks stabilised intermediates selectively^31,32,45^.

HFM1 showed strong enrichment at slow-resolving hotspots but little signal at fast-resolving sites in both B6 and hybrid mice (Figure 2C, Figure S2G,I). Notably, the difference between hotspot classes is larger for HFM1 than for BLM (Figure 2C, Figure S2I), indicating that slow-resolving intermediates are not only longer-lived but also more likely to engage ZMM-associated stabilisation.

Because HFM1 acts early in the ZMM pathway, these observations suggest an early decision to repair breaks in fast-resolving genes with non-crossover. To test this directly, we generated an *Mlh3*^-/-^ mouse, in which almost all crossovers are lost and strand-exchange intermediates that would otherwise form crossovers are repaired as non-crossovers^47^. Strikingly, the separation of hotspots into fast- and slow-resolving classes in *Mlh3*^-/-^ testes is similar to that in wild-type (Figure 2D), indicating that choice of fast versus slow repair mode is established upstream of MLH3-dependent crossover designation.

Together, these observations indicate that fast-resolving hotspots follow a pathway in which strand-exchange intermediates are short-lived and committed to non-crossovers early, whereas slow-resolving hotspots form intermediates that persist long enough to acquire stabilising factors needed for crossover formation.

### Gene expression in early meiosis predicts rapid resolution

The separation of hotspots into fast- and slow-resolving classes suggests the existence of a specific genomic feature that distinguishes them. Because many genomic features are correlated, an enrichment alone would not establish causality; instead, the relevant feature should *predict* the class to which each hotspot belongs. We therefore evaluated a broad set of genomic variables using single-feature statistical classifiers to determine which, if any, could accurately predict fast- and slow-resolving hotspots.

Features previously implicated in recombination behaviour, namely replication timing, GC-content, transposable-element density, and distance to telomeres, were poor predictors (Figure 3A, Figure S3A-F). Strikingly, however, fast-resolving hotspots were almost exclusively located within gene bodies: 95% lie between transcription start and end sites, whereas slow-resolving hotspots exhibited a distribution similar to the genomic background (47% in genes; Figure 3B). This repair behaviour was highly local and gene-specific, with adjacent genes often harbouring hotspots of opposite classes (Figure S3G), suggesting that the relevant information is encoded at the level of individual genes rather than broad chromosomal domains.

**Figure 3.**
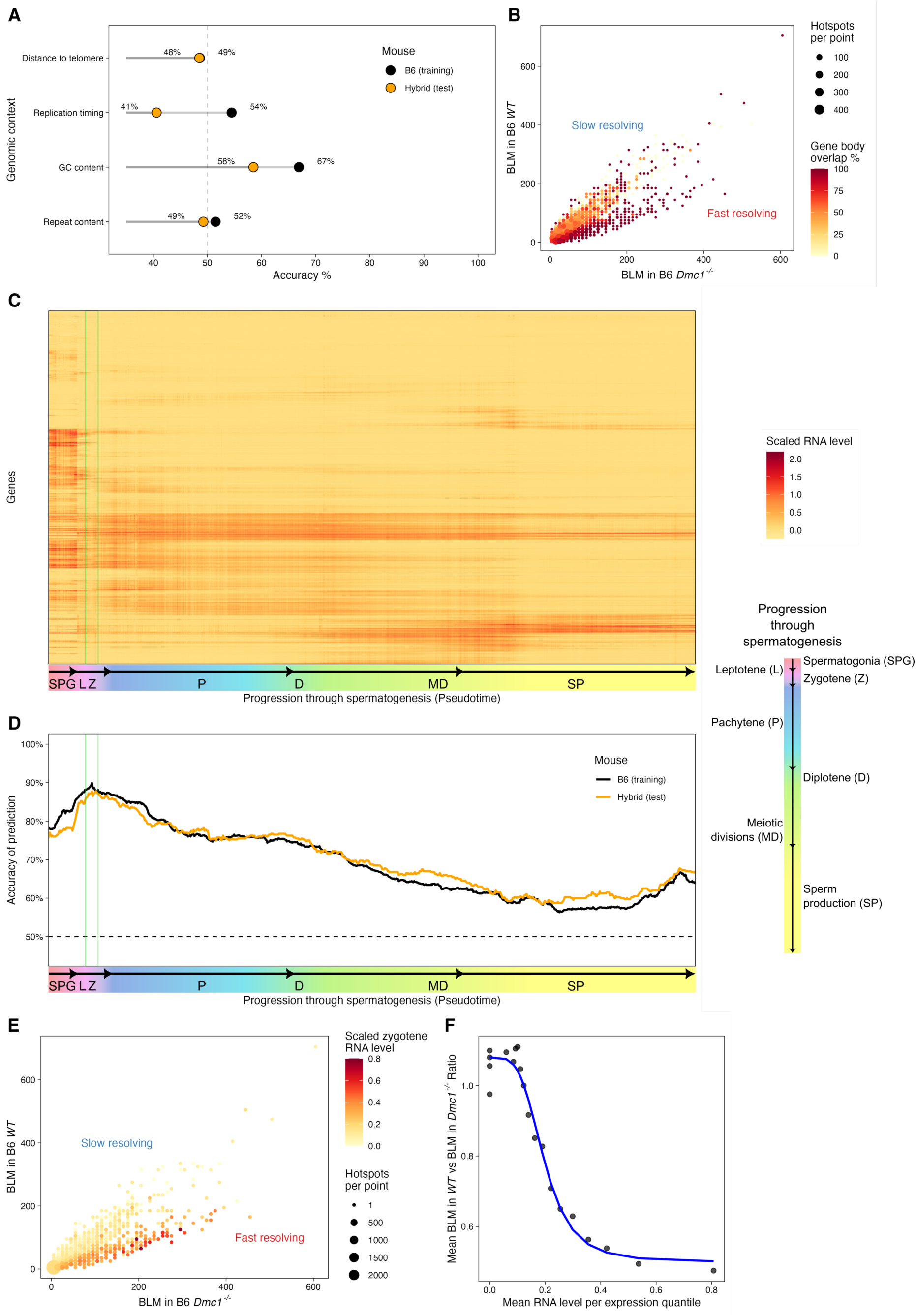
Gene expression in Zygotene predicts rapid strand-exchange resolution across mouse sub-species. **(A)** Prediction accuracy of individual genomic features for classifying hotspots as fast- or slow-resolving. Logistic regression models (STAR methods) were trained on B6 hotspots (black) and tested on hybrid hotspots (orange). The dashed line indicates random classification (50%). See Figure S3 for a more detailed assessment. **(B)** Cells with fast-resolving hotspots are strongly enriched for overlap with genes. Scatterplot of BLM signal in wild-type B6 relative to *Dmc1^-/-^* mice, with hotspots aggregated into cells of a 75x75 grid (each point represents one grid cell). Point size reflects the number of hotspots per cell; colour indicates the proportion of hotspots overlapping gene bodies. **(C)** Heatmap of single-cell RNA-seq levels across spermatogenesis in B6 mice. Each row represents a gene; columns represent pseudotime intervals across meiosis. RNA abundance in each time window was used to predict hotspot class; the zygotene window (green lines) is highlighted (Data from^48^). **(D)** Prediction accuracy of RNA level for classifying genic hotspots as fast- or slow-resolving across time windows in meiosis. Four distinct metrics (STAR Methods) consistently identify zygotene expression as the most informative stage. **(E)** As in (B), but for RNA level of genes overlapping hotspots. Grid cells are coloured by the mean log-scaled zygotene RNA level of genes harbouring the hotspots, illustrating concordance between zygotene expression and rapid strand-exchange resolution. **(F)** Relationship between strand-exchange–dependent BLM signal (WT/*Dmc1^-/-^* ratio) and zygotene RNA level for genic hotspots. The sigmoidal Hill-type fit^49^ (blue curve), indicates a threshold-like association between RNA abundance and intermediate lifespan.

A natural hypothesis is that a feature linked to transcription of genes distinguishes fast-resolving hotspots from other hotspots. Spermatogenesis involves a highly choreographed transcriptional programme in which thousands of genes are activated at distinct developmental stages (Figure 3C)^16-19^. If a transcription-associated feature determines how strand-exchange intermediates are handled, then genes expressed at the key developmental stage should preferentially harbour fast- or slow-resolving hotspots. To identify this stage, we leveraged single-cell RNA-sequencing data across ∼1,000 narrow pseudotime intervals spanning meiosis in B6 mice^48^. We used the RNA level of each gene within each time interval to predict whether a hotspot overlapping that gene would be fast- or slow-resolving (STAR Methods).

This analysis identified the gene expression pattern in zygotene, the early meiotic stage when homologous chromosomes undergo pairing, as the strongest predictor of rapid hotspot resolution, distinguishing fast- and slow-resolving hotspots with high accuracy (AUC = 0.96; accuracy = 90%) (Figure 3D-E, Figure S3F). The gene expression-repair relationship was highly non-linear: the average persistence of strand-exchange intermediates dropped sharply over a narrow range of RNA levels, consistent with a switch-like rather than gradual transition between fast- and slow-resolving behaviour (Figure 3F).

To assess whether this relationship generalises across mouse sub-species and *Prdm9* alleles, we used B6 zygotene RNA levels to estimate, for every mouse gene, how breaks *would* be expected to resolve if a hotspot were to occur within that gene. This yielded 4,470 out of 19,262 spermatogenesis genes predicted to support fast resolution. Testing these predictions in B6×CAST hybrids showed that, despite distinct PRDM9 specificities marking an almost entirely different set of hotspots, zygotene expression in B6 predicts hotspot class in the hybrid with high accuracy (88%), including for genes that do not harbour hotspots in B6 (84%).

This finding clarifies the direction of causality. Because predictions made in B6 extend to other sub-species at sites that are not broken in B6, this establishes that the transcriptional features that predict rapid resolution are not a consequence of breaks themselves. We conclude that the signal reflects a transcription-linked feature of genes with high RNA levels in zygotene, consistent with repair-pathway divergence being established at this early stage for them. We refer to these genes as fast-resolving genes henceforth.

### Structural and epigenetic features of prior transcription define micro-domains of rapid repair

As cells enter meiosis and DNA breaks are formed, transcription drops sharply due to global pausing of RNA Polymerase II (Pol II), with only a small number of genes escaping silencing^50-53^. Transcription restarts in pachytene, even though numerous recombination intermediates remain to be repaired, with ∼6,000-8,000 genes becoming reactivated^50,51^. Paradoxically, however, repair mode is best predicted by the RNA signature of zygotene, when transcription is maximally repressed (Fig. 3D).

The RNA produced before meiotic silencing is unusually stable and zygotene RNA levels largely reflect prior transcriptional activity^50^. As a conservative control, we nevertheless tested whether active transcription at a low level might contribute to rapid resolution. Pol II engages a specific DNA strand during transcription; therefore, a conflict between transcription and recombination machineries is expected to produce strand-biased perturbations of strand-exchange intermediates^7^. However, we did not observe any bias between the transcribed and non-transcribed strands (Figure S4A). Likewise, R-loops (non-canonical DNA:RNA structures), which can impact repair in somatic cells^54^, showed no difference between fast- and slow-resolving hotspots (Figure S4B). These results do not support a direct interaction between active transcription and strand-exchange intermediates, consistent with minimal transcription at this stage.

We therefore hypothesised that the effect arises from a persistent feature established during earlier transcription, such as transcription-associated histone modifications or structural changes in genome organisation. Among 42 chromatin-associated features examined, H3K36me3, a hallmark of transcriptional elongation deposited co-transcriptionally by SETD2, was the strongest predictor, correctly classifying 94% of hotspots in B6 and 90% in the hybrid (Figure 4A-B, Figure S4C,G). H3K36me3 persists along gene bodies after transcription ceases, preserving a footprint of prior activity across the transcribed unit. Other transcriptional elongation-associated chromatin marks showed comparable predictive performance (Figure 4A, Figure S4C,G), reinforcing the idea that features of prior transcriptional activity distinguish fast- and slow-resolving hotspots. Meiotic stage-resolved profiles identified mid-zygotene as the window when H3K36me3 best distinguished hotspot classes (Figure 4C, Figure S4D), consistent with the predictions based on gene expression (Figure 3D).

**Figure 4.**
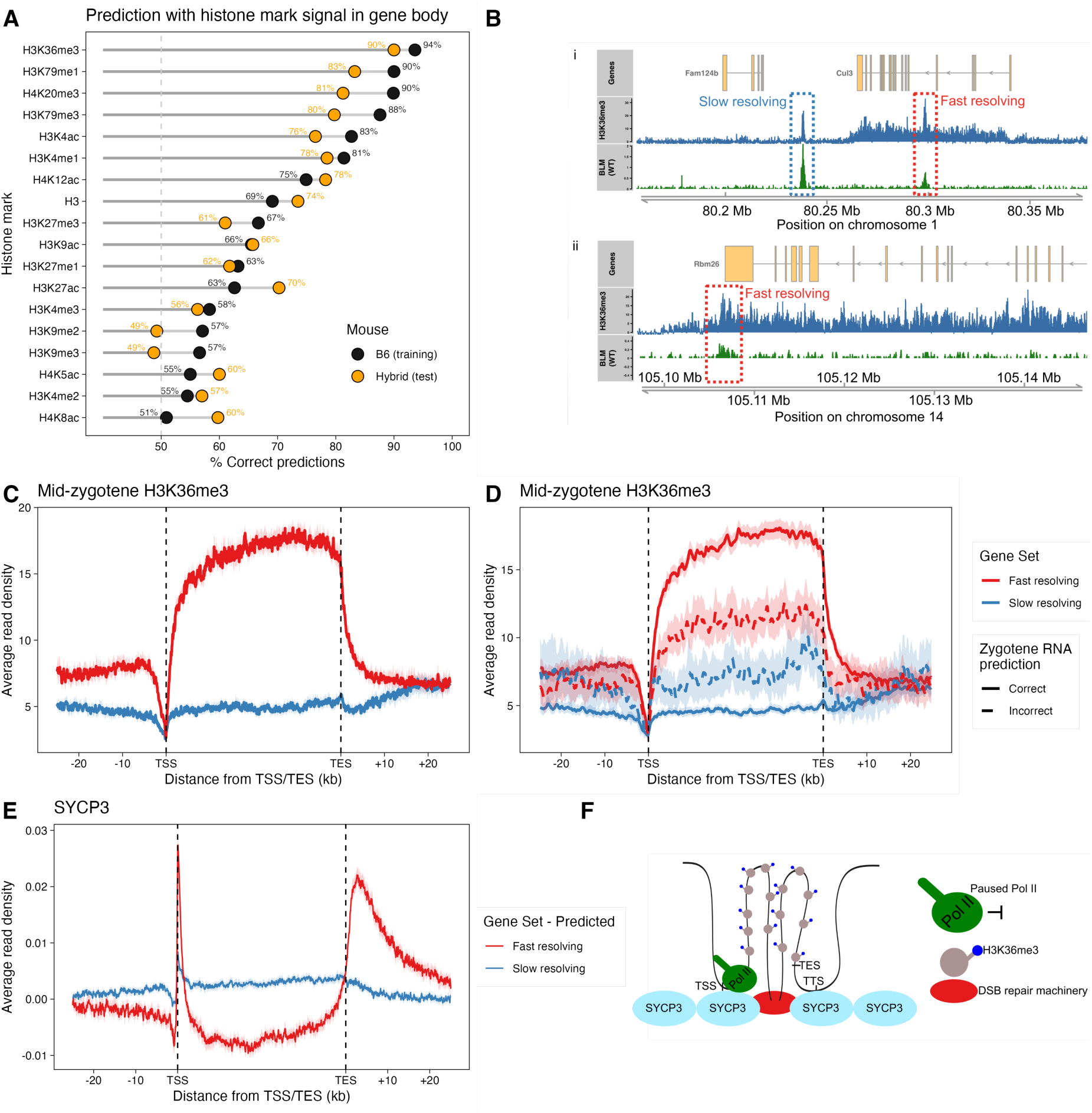
Chromatin and structural features of prior transcription define micro-domains of rapid strand-exchange resolution. **(A)** Prediction accuracy of gene body chromatin marks for classifying hotspots as fast- or slow-resolving. Logistic regression models (STAR methods) were trained on B6 hotspots (black) and tested on hybrid hotspots (orange). H3K36me3 and other transcription-associated features strongly predict rapid strand-exchange resolution. **(B)** Representative examples of B6 fast- and slow-resolving hotspots with H3K36me3 ChIP-seq (blue) and BLM ChIP-SSDS (green). Although H3K36me3 is locally deposited at hotspots by PRDM9, fast-resolving hotspots are embedded within broader H3K36me3-enriched transcriptional domains, whereas slow-resolving hotspots lie outside such domains. (i) A slow-resolving hotspot located outside transcriptionally marked regions contrasted with a fast-resolving hotspot within the *Cul3* transcription domain (ii) A fast-resolving hotspot located downstream of the annotated *Rbm26* transcript end site (TES), yet within an extended H3K36me3 domain reflecting continued RNA Polymerase II (Pol II) activity beyond the canonical TES. **(C)** Metagene profiles of mid-zygotene H3K36me3 across genes rescaled to a common length (40 kb, 50 bp smoothing; data from^56^). Genes harbouring fast-resolving hotspots exhibit elevated H3K36me3 across gene bodies relative to those harbouring slow-resolving hotspots. (TSS, transcription start site). **(D)** Elevated H3K36me3 levels are found in genes with fast-resolving hotspots even among genes with low RNA expression in zygotene, indicating that chromatin features of prior transcription are more closely associated with repair fate than RNA abundance. Metagene profiles of mid-zygotene H3K36me3 are shown for genes harbouring fast- (red) or slow- (blue) resolving hotspots in B6 or hybrid mice. Solid lines denote genes correctly classified by zygotene RNA level; dashed lines denote misclassified genes. **(E)** Metagene profiles of SYCP3, a core component of the meiotic chromosome axis in genes predicted to be fast-(red) or slow- (blue) resolving (data from^55^). Genes with fast-resolving hotspots show enhanced axis association at both ends, consistent with gene loops. **(F)** Model summarising the proposed regulatory architecture. Genes harbouring fast-resolving hotspots form H3K36me3-enriched chromatin micro-domains with axis tethering at both ends, biasing strand-exchange intermediates toward rapid, predominantly non-crossover repair. Breaks outside these domains persist longer and are more likely to contribute to crossover formation (TTS, transcription termination site, where Pol II disengages from the template).

Strikingly, an H3K36me3 signal encompassed the small minority of fast-resolving hotspots located outside genes (Figure 4B). These hotspots localised immediately downstream of transcript end sites within H3K36me3-retaining regions. Consistent with this, high H3K36me3 signal was also observed in the minority of fast-resolving hotspots located in genes with low zygotene RNA level (Figure 4D, Figure S4E).

Meiotic chromosomes are organised into DNA loop-axis structures, and the axis preferentially engages regulatory elements such as promoters and insulator sites^55^. Because fast-resolving hotspots are associated with genes retaining chromatin marks from prior transcription, we asked whether their gene boundaries also show distinctive axis engagement. Strikingly, genes predicted to be fast-resolving displayed strong enrichment for axis association at both their transcription start and termination sites (Figure 4E). In contrast, predicted slow-resolving genes showed only a weak promoter signal and no detectable enrichment at termination sites (Figure 4E).

These differences in boundary-axis contacts indicate that fast- and slow-resolving genes occupy distinct structural and chromatin configurations during meiosis. Dual anchoring at both gene ends could position recombination intermediates within more constrained structural domains in which D-loop extension or stabilisation of factors such as HFM1 is limited (Figure 4F). Recent studies in budding yeast show that axis proteins stabilise crossover-designated recombination intermediates, underscoring how chromosome architecture can directly influence repair-pathway choice^57,58^. Other contributors, such as transcription-linked chromatin marks or paused RNA Polymerase II at promoters, could also impact local repair dynamics. Together, these differences provide a structural context for the rapid and predominantly non-crossover repair at fast-resolving hotspots.

Although dozens of breaks persist into pachytene, and crossover precursors (chiasmata) persist further into diplotene, their persistence in genes with high expression in these later stages is strongly suppressed (Figure S4F). The structural and chromatin architecture of fast-resolving genes (Figure 4F) may therefore explain how transcription can resume safely while recombination remains active.

### Cross-species conservation of crossover suppression in fast-resolving genes

In humans, crossovers are known to occur less frequently within genes than expected from the genomic average, including in genes expressed during spermatogenesis^12,14^. We asked whether this phenomenon can be explained by the regulatory principle revealed here in mice. Specifically, if transcriptionally-marked genes in zygotene are channelled into rapid, non-crossover repair, then crossover suppression should be specific to this subset. Because the spermatogenic programme is largely conserved between mouse and human^19^, we identified orthologues of predicted fast-resolving mouse genes in humans (∼4,500) and compared both present-day and ancestral crossover patterns at these loci.

In a large human pedigree^14^, crossover frequency was ∼5-fold reduced within predicted fast-resolving genes (p=7.8x10^-233^), whereas no reduction was observed in other spermatogenesis genes relative to remaining genes (Figure 5A). Because fast-resolving genes were defined based on transcriptional features in male meiosis, we asked whether the same gene set also exhibits crossover suppression in females. While crossover number and localisation differ markedly between the sexes in general^14^, we observed a similar degree of suppression in females (Figure 5B). These effects are substantially greater than the ∼2-fold suppression reported previously across spermatogenesis genes^14^. Together, these results show that crossover suppression in humans is concentrated specifically in genes transcriptionally-marked in zygotene rather than spermatogenesis genes at large.

**Figure 5.**
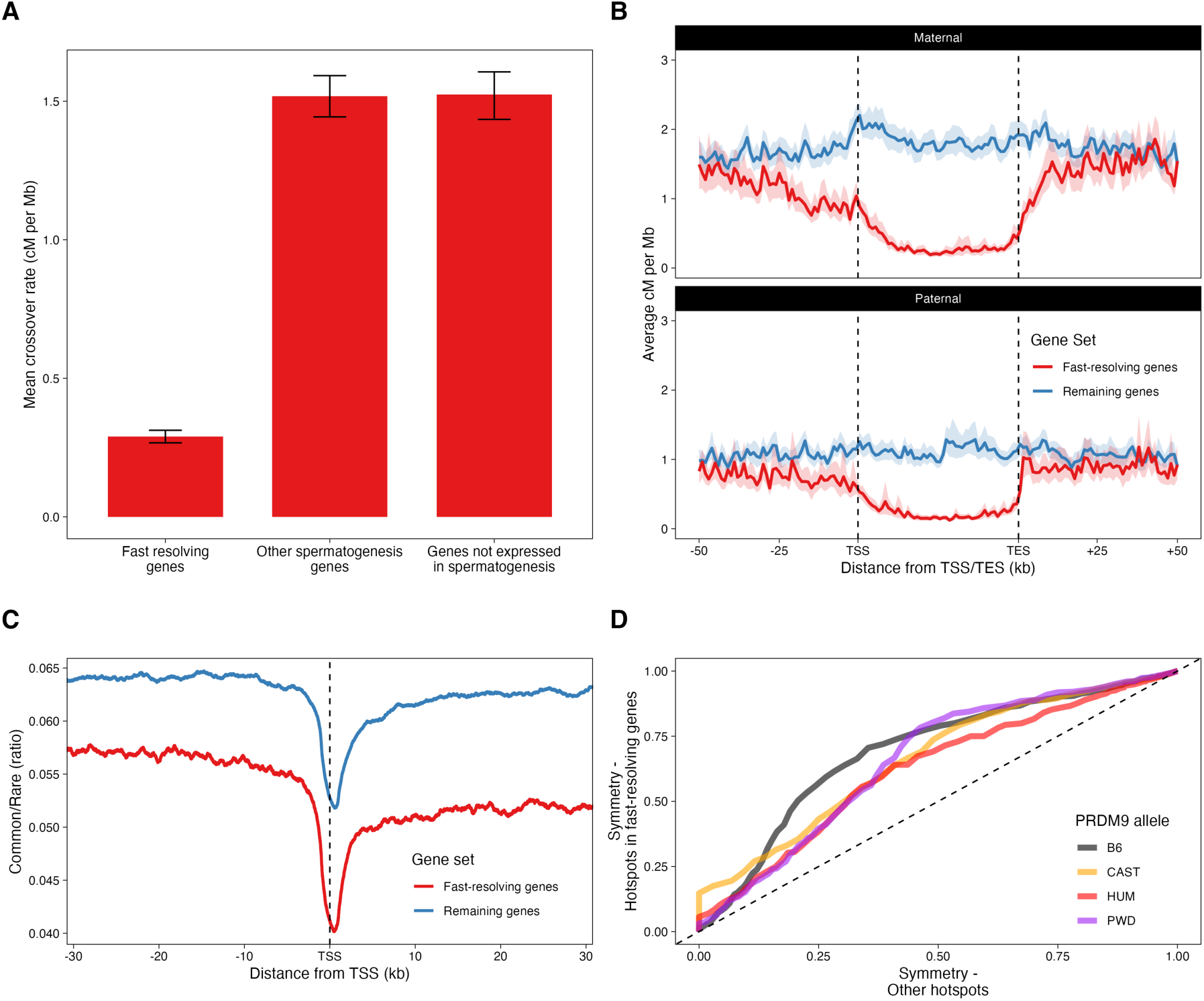
Conserved crossover suppression in humans and increased PRDM9 binding symmetry in fast-resolving genes. **(A)** Sex-averaged crossover frequency within gene bodies of human orthologues of predicted fast-resolving genes (n=4,227) compared with other spermatogenesis-expressed protein-coding genes (n=7,152) and remaining human protein-coding genes (n=6,524). Fast-resolving gene orthologues exhibit significantly reduced crossover frequency relative to other genes. **(B)** Human maternal (top) and paternal (bottom) crossover-rate metagene profiles for orthologues of predicted fast-resolving genes (red) versus all other protein-coding genes (blue). Gene bodies were scaled to 40 kb, and plotted using 1 kb bins. Crossover suppression within fast-resolving gene orthologues is evident in both sexes. **(C)** Ratio of common (minor allele frequency > 10^-2^) to rare (minor allele frequency < 10^-4^) single-nucleotide variants in human populations (gnomAD), plotted relative to transcription start sites of predicted fast-resolving genes (red) and other genes (blue), using a 2 kb-rolling smoothing window. Reduced common-to-rare variant ratios near fast-resolving genes are consistent with higher functional constraint and decreased crossing over. **(D)** Quantile-quantile plot of PRDM9 binding symmetry at hotspots activated by different PRDM9 alleles in mouse crosses, B6xPWD, B6^HUM^xPWD, B6^HUM^xCAST (B6^HUM^ denotes B6 mice carrying two copies of the *Prdm9*^HUM^ allele). Both wild-type and PRDM9^HUM^ hotspots exhibit significantly greater PRDM9 binding symmetry at predicted fast-resolving hotspots that at other hotspots (Bonferroni corrected p-values: B6, 1.2x10^-11^; CAST, 7.6x10^-15^; HUM, 7.7x10^-7^; PWD, 3.2x10^-7^). Increased symmetry promotes efficient homologue engagement and pairing^11^, suggesting that breaks within transcriptionally critical genes may provide a selective benefit despite crossover suppression.

In linkage-disequilibrium (LD)-based maps capturing ancestral recombination events over thousands of generations, the same suppression pattern was evident in diverse human populations (Figure S5A). The pattern also held in cattle, where both male and female recombination maps^59^ revealed reduced crossover density in orthologues of predicted fast-resolving genes (Figure S5B). Thus, crossover suppression in this gene set is conserved between the sexes and across widely-diverged mammalian lineages spanning at least ∼90 million years of evolution since their last common ancestor.

Predicted fast-resolving genes are highly conserved and enriched for essential processes including DNA repair, chromatin organisation, and RNA metabolism (Figure S5C). Functional annotations exhibited strong overrepresentation of housekeeping functions and higher haploinsufficiency scores than other genes (Figure S5D), indicating that these dosage-sensitive loci are disproportionately subject to rapid repair.

Although the average mutation rate in these genes is comparable to other genes genome-wide (Figure S5E), genetic variation is strongly depleted in them in both humans and mice. In humans, fast-resolving genes exhibit ∼20% reduction in the relative abundance of common versus rare variants (Figure 5C, Figure S5F-G), and in mouse they show 20% lower inter-subspecies divergence (B6 vs CAST). These patterns are consistent with strong purifying selection and loss of linked diversity in recombination-suppressed regions^2^. Consistent with reduced genetic diversity, hotspots in these genes are significantly more likely to have PRDM9 bound symmetrically on both homologues than hotspots elsewhere in the genome (Figure 5D). This suggests that they provide especially effective sites for homologue pairing and synapsis. Together, these findings reveal a conserved class of critical, dosage-sensitive genes with rapid repair and suppressed crossing over across species.

## Discussion

Meiosis must execute two demanding tasks in parallel: repairing hundreds of programmed DNA double-strand breaks (DSBs) and coordinating a transcriptional programme essential for germ cell development. This is inherently paradoxical. In early meiosis, when breaks are formed and homologues begin to pair, global repression of RNA Polymerase II stalls most transcription, and cellular activity is sustained largely by stabilised transcripts from earlier transcription^50,51^. Yet transcription resumes in pachytene while many recombination intermediates remain unresolved, and crossover precursors—multi-protein–DNA joint molecules called chiasmata—persist further into diplotene^1^. Because meiotic breaks are enriched within gene bodies in many mammals, including mouse and human^10,11^, this creates a fundamental risk: if chromatids carrying genes required for later developmental stages are occupied by repair intermediates or future chiasmata, timely transcription could be impeded.

In males, this conflict is acute when thousands of genes are reactivated in pachytene^51^. In females, meiosis introduces an additional challenge: recombination initiates in the foetal ovary, but oocytes complete meiosis decades later after transcriptional reactivation and dramatic cellular growth^60,61^. Avoiding crossovers within genes that must remain transcriptionally accessible at this stage may be especially important in the female germline. How meiotic cells mitigate these risks in both sexes has remained unclear.

Here we identify a regulatory principle that resolves this long-standing paradox. Across hotspots genome-wide, meiotic DSBs segregate into two kinetically distinct classes that diverge early in prophase. Fast-resolving breaks form strand-exchange intermediates that are short-lived on average and are overwhelmingly repaired as non-crossovers. Slow-resolving breaks persist long enough to be stabilised, and a subset of them ultimately form nearly all crossovers. While most genes are expressed at some point in the testis, fast-resolving hotspots occur almost exclusively within a specific, deeply conserved subset of ∼4,500 genes enriched for key nuclear functions.

Crucially, transcriptional and chromatin features in B6 males predict repair behaviour across mouse sub-species, sexes, and across widely-diverged mammalian lineages. Because these predictions apply to breaks that are absent from the original genetic background, the signal cannot arise as a consequence of hotspot activity itself. Instead, it reflects gene-intrinsic properties that are established independently of meiotic breaks and influence their repair once they occur. As a result, whenever recombination occurs within these loci—regardless of which PRDM9 allele creates the hotspot—breaks are expected to resolve rapidly and predominantly as non-crossovers. Transcriptional marking of genes therefore provides the strongest known basis for how meiotic cells bias recombination toward crossover or non-crossover repair across the genome.

Although perturbation experiments will ultimately be required to establish molecular causality, the ability to predict repair fate across genomes with distinct hotspot landscapes places strong constraints on the underlying mechanism. Gene expression patterns and transcription-associated chromatin marks, such as H3K36me3 in gene bodies, point to zygotene as the critical stage, when homologues are pairing but not yet stably synapsed. In yeast, recent evidence has suggested that repair intermediates can be repositioned through direct interaction with the transcriptional machinery^7,62^. However, increased non-crossover repair in fast-resolving genes and the absence of any compensatory increase in crossovers in their flanking regions indicate that a distinct regulatory mechanism operates in mammals. H3K36me3 has previously been implicated in DSB repair pathway choice in somatic cells, where it promotes homologous recombination over end-joining repair^63-65^. In contrast, our data reveal a previously unrecognised layer of regulation in meiosis, in which transcription-associated loci regulate non-crossover versus crossover outcomes within homologous recombination. In this context, dual axis tethering at transcription start and termination sites of fast-resolving genes may create constrained chromatin micro-domains in which D-loop extension or engagement with stabilising factors is limited, favouring rapid dissolution into non-crossovers (See Supplementary Text for further discussion of underlying mechanism).

This framework reconciles previously puzzling observations. First, it explains how transcription can resume safely in pachytene despite the presence of unrepaired breaks: breaks in transcriptionally critical genes are on a rapid, early-resolution timescale that reduces the likelihood of conflict when transcription resumes. Second, it accommodates the dual demands of crossover assurance and interference by allowing breaks elsewhere to persist long enough to promote optimal number, spacing, and placement of crossovers.

The evolutionary consequences of this regulatory strategy are substantial. Genes hosting fast-resolving hotspots form a conserved class of ∼4,500 genes enriched for housekeeping, chromatin, and RNA-processing functions. These genes have strongly reduced diversity despite normal mutation rates, consistent with strong functional constraint and loss of linked genetic variation in crossover-suppressed regions. Although reduced crossing over is predicted to decrease the efficacy of natural selection in these regions, it may also preserve functionally constrained haplotypes. An immediate implication is a limit on how precisely disease-causing variants in these genes can be fine-mapped. Because crossover suppression in them is evolutionarily conserved, this limitation is likely to persist even as increasingly diverse populations are studied.

Why, then, do meiotic breaks occur within such critical genes? Our results suggest that, within the constraints of PRDM9-driven break placement, this localisation has functional consequences. Genes within this conserved set exhibit substantially more symmetrical PRDM9 binding between homologues, a feature that is predicted to facilitate efficient homologue engagement and synapsis. In species lacking PRDM9, selection on promoter architecture may enforce comparable symmetry^66^. Determining how recombination partitioning operates in species with promoter-driven hotspots (e.g., birds, many teleosts^67,68^) or in mammals with atypical recombination landscapes (e.g., canids^69^) will illuminate how ancient and widespread this regulatory mechanism is.

We therefore propose a principle of partitioned risk in meiotic recombination. DSBs in transcriptionally important genes are tolerated because strand-exchange intermediates in them are directed toward rapid, predominantly non-crossover repair, preserving transcriptional access. Breaks elsewhere persist and are stabilised to generate crossovers that ensure faithful chromosome segregation. Meiotic cells thus coordinate gene expression with recombination outcomes, enabling a balance between genome stability, transcriptional integrity, and evolutionary constraint.

## STAR Methods

### Mouse breeding and husbandry

All animal procedures were performed in accordance with UK Home Office Animal (Scientific Procedures) Act 1986, with procedures reviewed by the clinical medicine animal welfare and ethical review body at the University of Oxford and conducted under project license PAA2AAE49. Animals were housed in individually ventilated cages, provided with food and water *ad libitum*, and maintained on a 12-h/12-h light–dark cycle (150–200 lux). The only reported positives on FELASA health screening over the entire time course of these studies were for *Entamoeba spp*. Experimental groups were determined by genotype and were therefore not randomised, with no animals excluded from the analysis. Sample sizes for ChIP analysis were selected on the basis of previously published studies^28^, and all phenotypic characterisation was performed blind to experimental group. Genetically modified mice harbouring the human PRDM9 B allele (*Prdm9^tm1.1(PRDM9)Wthg^*) were generated inhouse^11^. CAST/EiJ mice were provided by Jonathon Godwin at the Sir William Dunn School of Pathology, University of Oxford. C57BL/6J mice were purchased from Charles River Laboratories.

### Mouse genetic modifications

Genetic ablation of *Dmc1* and *Mlh3* was performed as previously described, analysing founder knock-out mice^43^. Briefly, B6CASTF1 zygotes, generated from superovulated C57BL/6J females homozygous for the *Prdm9^tm1.1(PRDM9)Wthg^* allele mated with wild-type CAST/EiJ males, or wild-type C57BL/6J zygotes were electroporated with 65 ng/μL of two sgRNAs per target gene (see Table S4) and 650 ng/μL NLS-Cas9 protein in OptiMem buffer (2x square wave 30V pulses, 3 ms duration and 100 ms interval, GenePulser X-cell (BioRad)). Embryos were cultured overnight to the 2-cell stage and surgically transferred into psuedopregnant CD1 females. Resulting founder offspring were genotyped using PCR amplicons spanning the CRISPR-Cas9 target sites (Table S4), followed by Sanger Sequencing. Male founder offspring harbouring biallelic loss-of-function mutations (frame-shift or *in cis* deletion between the two CRISPR sites) were selected for analysis. A functional knock-out of *Dmc1* was confirmed in meiotic spreads by immunohistochemistry using a rabbit polyclonal anti-DMC1 antibody (Proteintech 13714-1-AP) to test absence of DMC1 foci (Table S5) as previously described^43^. Functional ablation of *Mlh3* was confirmed by immunohistochemistry using a mouse anti-MLH1 antibody (BD, 51-1327GR) to confirm a lack of crossover events (Table S5), as previously described^43^. Details of the mice used for the ChIP-SSDS analysis are shown in Table S5.

### BLM and HFM1 ChIP-seq maps

ChIP followed by SSDS was performed as described previously^39^ with some modifications^28^. The antibodies used were BLM (GeneTex GTX101303), RPA2 (Calbiochem RPA34-20) and HFM1 (GeneTex GTX65542). Sequencing was performed on an Illumina HiSeq X sequencer with 150-bp paired-end reads. The reads were trimmed to 75 bp before processing with the analytical pipeline to identify ssDNA^39^. Hotspots were called using our published peak-calling algorithm^11^. Tables S1-3 summarise the characteristics of the experimental data sets for BLM, RPA, and HFM1 ChIP-SSDS, respectively.

### Unsupervised clustering of hotspots and estimation of class membership of each hotspot

See Section ‘Statistical Framework for Unsupervised Clustering of Hotspots Into Two Classes’ in the Supplementary Text.

### Hotspot sets matched for DNA break frequency

Each hotspot with a high inferred probability of belonging to the fast-resolving (low BLM signal) class (p>0.75) was matched to a hotspot from the other class (p>0.75) with the most similar number of mapped SPO11-oligos^25^ within 500 bp of hotspot centres. Only pairs with <1% difference between their SPO11-oligo counts were considered valid matches.

### Signal aggregation across genomic intervals (metagene) plots

Coverage matrices were produced from bigWig files using the computeMatrix function of deepTools2^70^ For hotspots, the ‘reference point’ mode was employed. For genes, which are variable in size, the ‘scale regions’ mode was used. Coverage matrices were loaded into R version 4.4.1 (2024-06-14), mean coverage values calculated, and final figures plotted using ggplot2. 95% confidence intervals were generated for each plot using bootstrap resampling (n=1000).

### Relative frequency of non-crossovers in slow- and fast-resolving hotspots

Non-crossovers have short gene conversion tracts and can only be discovered if the homologous chromosomes have one or more sequence differences in that locus. Fast-resolving hotspots have fewer single-nucleotide polymorphisms (SNPs) on average, which reduces power to detect non-crossovers in them. To account for this difference, we estimated gene-conversion tract lengths in fast-and slow-resolving hotspots separately through a previously described approach^37^ that takes SNP density into account (Figure S2H). We used these inferred distributions to sample the locations of 100,000 hypothetical non-crossovers from hotspots in each set. A non-crossover was deemed to be ‘discoverable’ if it overlapped a SNP between B6 and CAST genomes^71-73^. We inferred that 10.5% of non-crossovers in fast-resolving and 16.9% in slow-resolving hotspots are discoverable. The total number of non-crossovers in each group was estimated by re-scaling the observed number of non-crossovers by their visibility.

### Classification and prediction of hotspots as fast- or slow-resolving based on genomic, transcriptomic, and epigenomic context

For classification and prediction using genomic context information, such as GC-content or replication timing, single-variable logistic classifiers were trained on B6 using the most confidently clustered hotspots in each class, scored by the inferred probability of belonging to either class.

Hotspots from in the hybrid mouse were used as test data. For subsequent transcriptomic/epigenomic classification models, the top B6 hotspots overlapping unique genes were used as training data and the hybrid hotspots overlapping unique genes were used as test data. To minimise noise in classification, we set a minimum threshold such that no hotspot across any training or test set had probability less than 50% of belonging to its assigned class. Equal number of hotspots from each class were used to prevent bias in training, and across models to maximise comparability, which was limited by the (smaller) fast-resolving class. Setting the threshold to 500 hotspots for each class satisfied these constraints in B6. In the hybrid, analyses were restricted to symmetric hotspots such that PRDM9 binding fraction *f* on each homologue is 0.4 < *f* <0.6. Consequently, a small threshold of 200 hotspots was required to satisfy the constraints.

For scRNA-seq data, models were generated using moving pseudotime windows of variable width to determine the most predictive time window. Genes present in the training and test datasets with no expression in spermatogenesis scRNA-seq were assigned an expression value of 0.

For chromatin marks datasets, classification models were again trained on B6 hotspots. Models were generated for the TSS region (-1000bp to +1000bp), and gene body region (from TSS + 1000bp to TES) for each gene. Average read densities for each region were quantified from publicly available bigwig/bedgraph files for each histone mark^56,74^ using the map function of BEDTOOLS. The effect of PRDM9 histone methylation was subtracted by removal of signal +/- 1000bp surrounding hotspots.

Model performance was assessed by accuracy of classification, Receiver-Operator Characteristic (ROC) Areas Under the Curve (AUC), Precision-Recall (PR) AUC, and by Matthews Correlation Coefficient^75^. Remarkably, each of these metrics identified the same zygotene window as the best performing in both training (B6) and test (hybrid) datasets.

### Non-linear Hill equation plots for scRNA-seq data

Zygotene-stage gene expression levels were quantified from single-cell RNA-seq data^48^ by averaging RNA counts in the pseudotime interval corresponding to the most predictive window identified above. These mean expression values were compared to BLM ChIP-SSDS signal in wild-type and *Dmc1^-/-^*backgrounds for hotspots overlapping these genes. A pseudo-count value of 1x10^-10^ was added to the expression values of every gene to ensure strict positivity. In the case of hotspots overlapping multiple genes, the gene with the highest RNA count in zygotene was retained. Genes were binned into 20 expression deciles based on their zygotene RNA levels, and the mean BLM ratio per bin was calculated as:

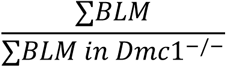

A non-linear Hill equation^49^ was fitted to the binned data using the nlsLM function from the minpack.lm package in R:

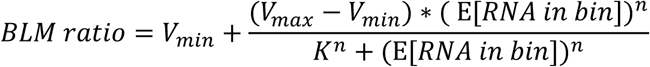

where *V_min_* and *V_max_* are the minimum and maximum plateau values, *K* is the half-maximal effective concentration, and *n* is the Hill coefficient controlling the slope. Parameter estimates and confidence intervals were derived from the model fit.

### Mouse gene sets from predictive modelling

For SYCP3 and H3K36me3 metagene profiles, fast-resolving genes were identified using the most predictive mouse scRNA-seq prediction model as protein-coding genes with score < 0.1 of belonging to the slow-resolving gene set (n=4,243). High-confidence slow-resolving genes were defined as protein coding genes with score > 0.9, including genes not present in the scRNA-seq expression matrix (n=10,355). Predicted gene lists were subsequently refined for downstream analysis using the mid-zygotene H3K36me3 level prediction model. Fast-resolving genes were defined as protein-coding genes with score < 0.1 (n=4,477).

### Gene ontology analysis

The g:GOSt functionality of g:Profiler^76^ was used for functional enrichment analysis of the predicted fast-resolving gene set above. Enrichment analysis was performed using default settings with the GO Biological Process data source, restricted to driver terms only.

### Gene ortholog selection and human / bovine gene annotations

Human and bovine orthologs of predicted gene sets were determined using the g:Orth functionality of g:Profiler^76^. For human and bovine analyses, fast-resolving genes were defined as the respective orthologs of the H3K36me3-defined fast-resolving gene set above, while non-fast-resolving genes were defined as all human or bovine protein-coding genes excluding orthologs of mouse genes with H3K36me3 predicted score < 0.5. Ortholog gene names were extracted from the output and intersected with the Gencode V47 human gene annotation and the UMD3.1.80 bovine gene annotation respectively, resulting in 4,227 (human) / 3,616 (bovine) fast-resolving gene orthologs; 13,676 (human) / 12,028 (bovine) remaining genes. For human analysis, we further defined non-fast-resolving spermatogenesis genes as the subset of the non-fast-resolving human gene set with orthologs in the mouse scRNA-seq data (n= 7,152), with genes not expressed in spermatogenesis defined as all remaining protein-coding genes.

### Crossover quantification from *Homo sapiens* and *Bos taurus* recombination maps

Pedigree-based crossover maps were obtained for humans and cattle from ^14^ and ^59^ respectively. Interval maps were converted into bigWig files with coverage defined as centiMorgans per Mb across each interval. Mean per-base scores per gene were determined using the bigWigAverageOverBed function from the UCSC genome browser analysis tools package Kent_tools/468-GCC-12.3.0.

### Human *de novo* mutation & variant allele datasets

De novo single nucleotide variants (DNMs) were collated from four publicly available whole-genome sequencing (WGS) trio studies^13,77-79^, yielding a total of 915,050 autosomal DNMs. Population single nucleotide variants were obtained from gnomAD v4.1^80^(WGS-only), and stratified into allele frequency (AF) bins using previously defined thresholds^10^. Common variants (AF 10^-1^–10^0^) comprised ∼5.3 million SNVs, and rare variants (AF 10⁻⁶–10⁻⁵) comprised ∼61.5 million SNVs

| PMID | Reference | Paper Title | DNM Count |
| --- | --- | --- | --- |
| 30545852 | <sup>77</sup> | Genome-wide de novo risk score implicates promoter variation in autism spectrum disorder | 233,954 |
| 27322544 | <sup>78</sup> | Parent-of-origin-specific signatures of de novo mutations | 35,752 |
| 31549960 | <sup>79</sup> | Large, three-generation human families reveal post-zygotic mosaicism and variability in germline mutation accumulation | 27,154 |
| 39843742 | <sup>13</sup> | Complete human recombination maps | 618,190 |

### De novo mutation and SNV density plots

To assess mutation density near gene features, counts of DNMs and gnomAD SNVs were calculated relative to distance from TSSs. For TSS analysis, distances between DNMs and TSSs within 40 kb were calculated, accounting for strand orientation. DNMs were binned into 1 bp intervals and aggregated by predicted repair category, yielding base-level DNM counts across regions flanking TSSs. These raw counts were normalized by the number of genes in each group (4,227 orthologs of fast-resolving genes and 13,676 for remaining genes). Fold enrichment at each position was computed as the normalized count at each bp distance divided by the mean normalized count in distal flanking regions (5-40 kb).

All enrichment plots were smoothed using a 2 kb moving average. Confidence intervals were generated using 100 bootstrap replicates, in which genes were resampled with replacement. Smoothed enrichment curves were re-calculated per replicate, and 95% confidence intervals were defined as the 2.5th and 97.5th percentiles at each base position. Population variant analyses followed the same procedure. gnomAD SNVs were stratified into rare and common AF categories based on allele frequency. Aggregation, normalisation, smoothing, and bootstrap-based confidence interval estimation were performed identically to the DNM analyses.

### Haploinsufficiency scores

Haploinsufficiency scores for human protein-coding genes were obtained from^81^ and intersected with the human orthologs of fast-resolving genes and remaining non-fast-resolving genes, leaving scores for 14,761 protein-coding genes (n=3,954 fast-resolving gene orthologs, n=10,807 other genes).

| Dataset | GEO Accession Numbers<br>(where available) | Reference(s) |
| --- | --- | --- |
| BLM ChIP-SSDS – B6, B6 *Dmc1*<sup>-/-</sup>,<br>hybrid, hybrid *Dmc1*<sup>-/-</sup> | This study |  |
| RPA ChIP-SSDS – B6 *Dmc1*<sup>-/-</sup>, hybrid<br>*Mlh3*<sup>-/-</sup> | This study |  |
| HFM1 ChIP-SSDS – B6, hybrid | This study |  |
| SPO11-linked oligos - B6 | GSE84689 | 25 |
| DMC1 ChIP-SSDS - B6 | GSE35498 | 82 |
| DMC1 ChIP-SSDS - Hybrid | GSE124991 | 36 |
| DMC1 ChIP-SSDS - PWDxB6,<br>PWDxB6<sup>HUM</sup> | GSE73833 | 11 |
| RPA ChIP-SSDS - B6, hybrid | GSE143582 | 28 |
| END-seq - B6, B6 *Dmc1*<sup>-/-</sup>, B6 *Hop2*<sup>-/-</sup> | GSE138915 | 42 |
| Crossover maps - Hybrid single-sperm<br>sequencing | GSE125326 | 36 |
| Crossover maps - Hybrid pedigree map |  | 37 |
| Non-crossover maps - Hybrid pedigree<br>map |  | 37 |
| Single-cell RNA-seq - B6 | GSE113293 | 48 |
| S9.6 CUT & TAG | GSE193401 | 83 |
| Histone marks ChIP-seq - B6 | GSE121760 | 74 |
| Stage-specific H3K36me3 - B6 | GSE132446 | 56 |
| SYCP3 ChIP-seq - B6, B6 *Spo11*<sup>-/-</sup> | GSE262343 | 55 |
| Crossover maps - Human maternal,<br>paternal, sex-averaged |  | 14 |
| Crossover maps - Cattle maternal,<br>paternal |  | 59 |
| Linkage disequilibrium maps - Human<br>Bagandan, European populations |  | 84 |

## Key Resources Table

Datasets used in the preparation of this manuscript. Mouse models generated in this study (*Dmc1^-/-^* and *Mlh3^-/-^*) are described in STAR methods and in Tables S4 and S5.

## Data and resource availability

We have generated the following whole-genome resources in this study:

- BLM ChIP-SSDS maps in B6 WT and *Dmc1^-/-^* and hybrid WT and *Dmc1^-/-^* (Table S1)
- RPA ChIP-SSDS maps in B6 *Dmc1^-/-^* and hybrid *Mlh3^-/-^* (Table S2)
- HFM1 ChIP-SSDS maps in B6 and hybrid WT (Table S3)

Data have been deposited in the NCBI Sequence Read Archive under BioProject accession **PRJNA1438681.**

## Acknowledgments

We thank Bernard de Massy and Simon Myers for helpful discussions. We are grateful to Jakob Weiss, Bernard de Massy, and Jordan Raff for critiquing the manuscript. This work was funded by the following grants:

Wellcome Trust grant 221761/Z/20/Z (AGH)

Wellcome Trust grant 095552/Z/11/Z (PD)

Wellcome Trust grants 090532/Z/09/Z and 20314/Z/16/Z (Core support to WHG)

## Supplementary Data and Supplementary Text

**Figure S1.**
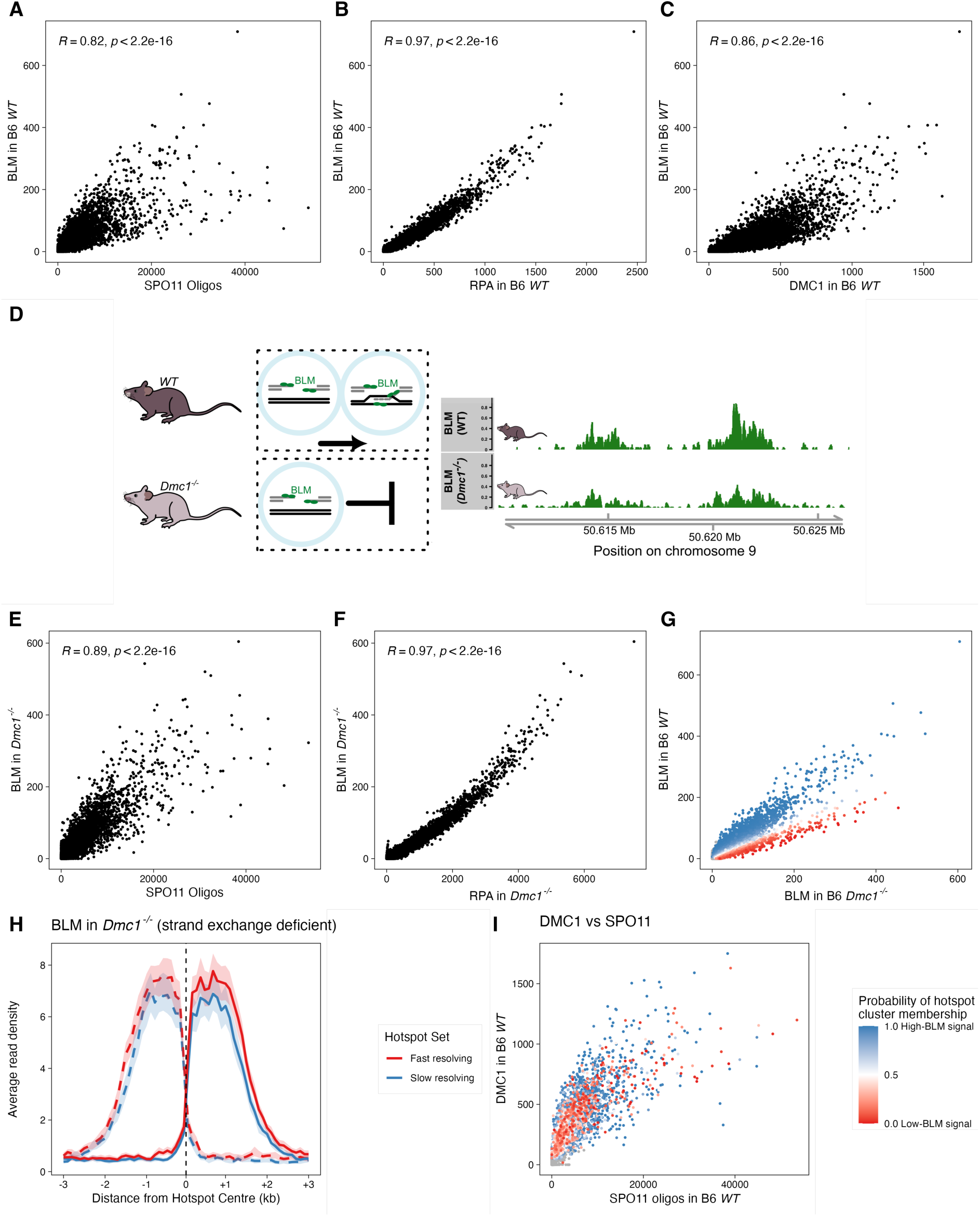
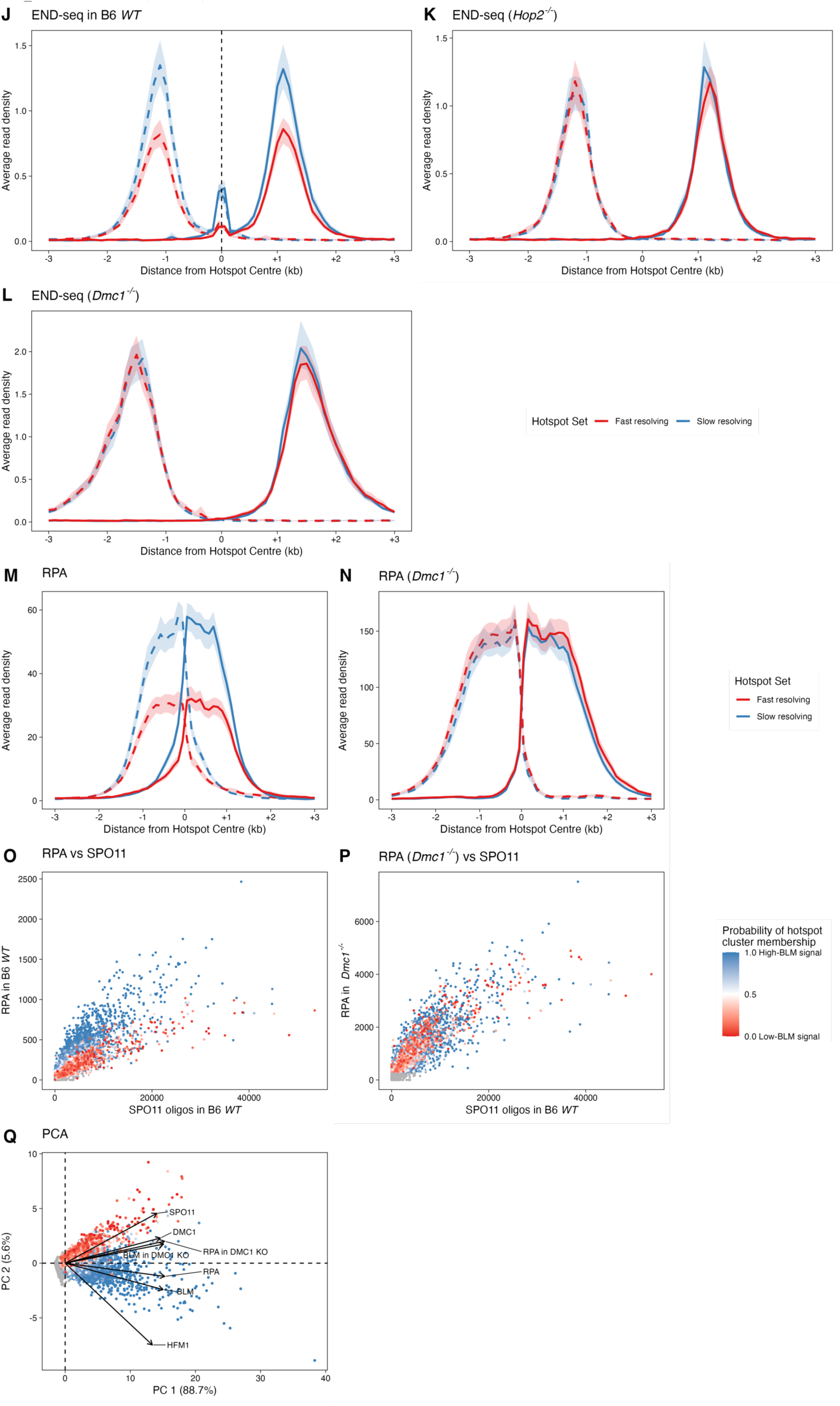
Two temporal modes of DNA repair revealed by genome-wide maps of recombination proteins. **(A)** Scatterplot of BLM ChIP-SSDS signal versus SPO11 oligo counts across 16,658 autosomal wild-type B6 hotspots (each point represents one hotspot), illustrating hotspot-to-hotspot variation in BLM signal relative to break frequency. **(B)** Scatterplot of BLM versus RPA ChIP-SSDS signal in wild-type B6 hotspots, showing close concordance between BLM and RPA, consistent with BLM interacting with RPA. **(C)** Scatterplot of BLM versus DMC1 ChIP-SSDS signal in wild-type B6 hotspots. Although binding of both proteins depends on resection, BLM exhibits substantial hotspot-to-hotspot variation relative to DMC1, consistent with association of BLM with strand-exchange intermediates and DMC1 with homology search intermediates. **(D)** (Left) Schematic illustrating that in *Dmc1*^-/-^ mice strand exchange does not occur; BLM ChIP-SSDS signal therefore reflects binding only to resected ssDNA intermediates. In wild-type (WT), BLM associates additionally with strand-exchange intermediates. (Right) Representative hotspots showing BLM ChIP-SSDS signal in WT and *Dmc1*^-/-^ mice. **(E)** As in (A) but for BLM in *Dmc1^-/-^* relative to SPO11 oligo counts, exhibiting higher correlation relative to BLM in wild-type. **(F)** Scatterplot of BLM versus RPA ChIP-SSDS signal in *Dmc1^-/-^*mice, showing close concordance. This high correlation is consistent with BLM and RPA binding the same resected-ssDNA substrate in the absence of strand exchange. **(G)** Scatterplot of BLM ChIP-SSDS signal in wild-type B6 versus *Dmc1^-/-^*, with hotspots coloured by the inferred probability of assignment to the high (blue) or low (red) signal classes. The plot shows the same class separation evident in Figures 1C–D: hotspots with elevated and reduced WT-specific BLM signal form distinct groups. **(H)** Average BLM signal in matched hotspots in *Dmc1*^-/-^ (100 bp smoothing, matching for DSB frequency as in Figure 1F) for the top (dashed) and bottom (solid) strands. No significant difference is observed between classes on average when strand-exchange intermediates are absent. **(I)** Scatterplot of DMC1 ChIP-SSDS signal versus SPO11 oligo counts, showing no separation between hotspot classes, consistent with comparable strand invasion efficiency. **(J)** END-seq in wild-type B6 hotspots shown for matched sets of fast- (red) and slow- (blue) resolving hotspots (matched for DSB frequency as in Figure 1F; data from^42^) for the top (solid) and bottom (dashed) strands. END-seq detects ssDNA–dsDNA junctions generated during DSB processing and strand exchange. Its characteristic profile reflects two components: (i) a central component where strand-exchange structures accumulate, and (ii) a flanking signal, which arises from a combination of resection endpoints^42^ and ssDNA in incomplete strand-exchange intermediates located outside the D-loop on the DSB-initiating chromosome^43^. In wild-type, slow-resolving hotspots show excess of both the central signal and the flanking signal, consistent with higher strand-exchange intermediate persistence. In strand-exchange mutants, the central component is absent and the flanking signal reflects resection-derived junctions alone. The similar END-seq profiles observed for fast- and slow-resolving hotspots in *Hop2^-/-^*and *Dmc1^-/-^* mice (panels K and L) provide further evidence that the divergence in wild type depends on strand-exchange intermediate persistence. **(K)** END-seq signal in *Hop2^-/-^* mice, shown for the same matched hotspot sets as in (J). This mutant lacks strand exchange; no separation between fast- and slow-resolving hotspots is observed, demonstrating that the divergence in WT END-seq signal in (J) is due to persistence of strand-exchange intermediates. **(L)** END-seq signal in *Dmc1^-/-^* mice, shown for the same matched hotspot sets as in (J). As for *Hop2^-/-^* mice, no separation between fast- and slow-resolving hotspots is observed in the absence of strand exchange. This again demonstrates that the divergence in WT END-seq signal in (J) is due to persistence of strand-exchange intermediates. **(M)** RPA ChIP-SSDS signal in wild-type B6, shown for matched sets of fast- (red) and slow-resolving (blue) hotspots (matching as in Fig. 1F) for the top (dashed) and bottom (solid) strands. RPA, which binds resected ssDNA, reproduces the class separation seen with BLM in wild type, supporting differences in ssDNA persistence at strand-exchange intermediates. **(N)** RPA ChIP-SSDS signal in *Dmc1^-/-^* mice, shown for the same matched hotspot sets as in (M). The difference disappears in *Dmc1^-/-^*, consistent with absence of strand-exchange intermediates. **(O)** Scatterplot of RPA ChIP-SSDS signal versus SPO11-oligo counts in WT mice across 16,658 autosomal hotspots (each point represents one hotspot). Points are coloured according to inferred class assignment as in (G) (high, blue; low, red). The plot shows the same class separation evident in Figures 1C–D, consistent with strand-exchange–dependent ssDNA persistence. **(P)** As in (O), but comparing RPA in *Dmc1^-/-^* to SPO11 oligos, confirming absence of strand-exchange–dependent separation. **(Q)** Principal Component Analysis (PCA) plot of key recombination measures across 16,658 autosomal hotspots. Points are coloured by inferred BLM cluster probability (high, blue; low, red). PCA separates hotspots according to strand-exchange–dependent behaviour, corroborating the clustering used in the main text.

**Figure S2.**
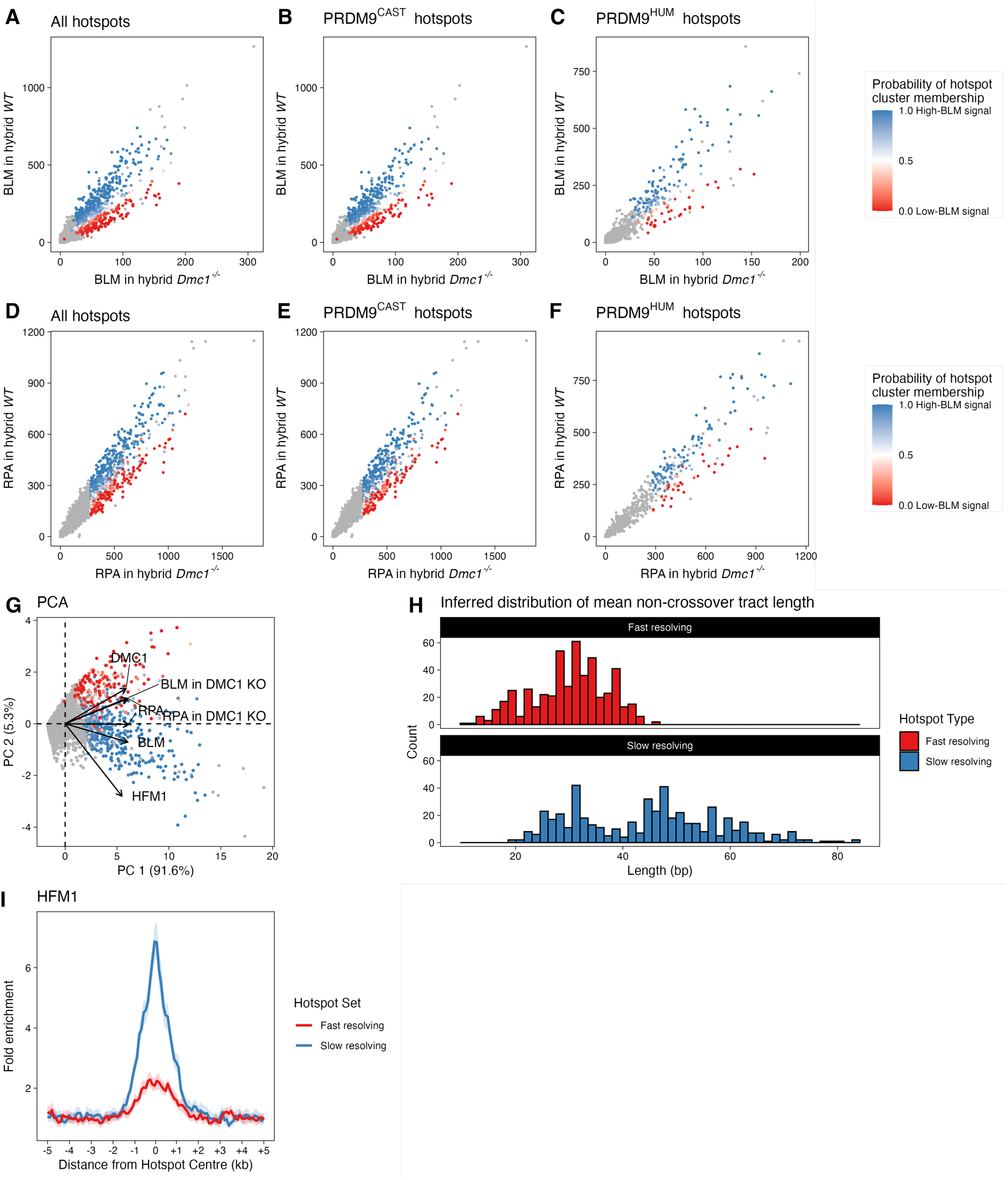
Fast and slow resolution generate distinct recombination outcomes. **(A)** Scatterplot of BLM ChIP-SSDS signal in wild-type hybrid relative to *Dmc1^-/-^* hybrid across 8,293 symmetric autosomal hotspots. Points are coloured by inferred probability of belonging to the slow-resolving (blue) or fast-resolving (red) cluster (as in Fig. 2). Hotspots with insufficient information for classification are shown in grey. Separation of hotspots observed in B6 is also observed in hybrid symmetric hotspots. **(B)** As in (A) but restricted to PRDM9^CAST^ hotspots (n=4,740), demonstrating that class separation is maintained in these hotspots. **(C)** As in (A) but restricted to PRDM9^HUM^ hotspots, showing similar separation and confirming allele independence of the separation. **(D-F)** As (A-C) but for RPA ChIP-SSDS signal. RPA reproduces the same strand-exchange–dependent class separation observed with BLM, confirming that divergence is not specific to BLM occupancy. **(G)** Principal Component Analysis (PCA) of key recombination measures across 8,293 symmetric autosomal hybrid hotspots. Colour scale reflects the inferred probability of each hotspot belonging to the slow-resolving (blue) or fast (red) clusters. Principal components separate hotspots according to strand-exchange–dependent behaviour, corroborating the cluster structure. **(H)** Fast- and slow-resolving hotspots differ systematically in single-nucleotide polymorphism (SNP) density, which impacts non-crossover detection. To correct for this bias, non-crossover tract lengths were explicitly modelled for each set (STAR Methods). The inferred mean tract length distributions are shown. **(I)** Average HFM1 ChIP-SSDS signal in matched sets of hybrid hotspots from slow-resolving (blue) and fast-resolving (red) clusters (matching and smoothing as in Fig. 2). Unlike fast-resolving hotspots, slow-resolving hotspots exhibit strongly elevated HFM1 association, consistent with stabilisation of strand-exchange intermediates.

**Figure S3.**
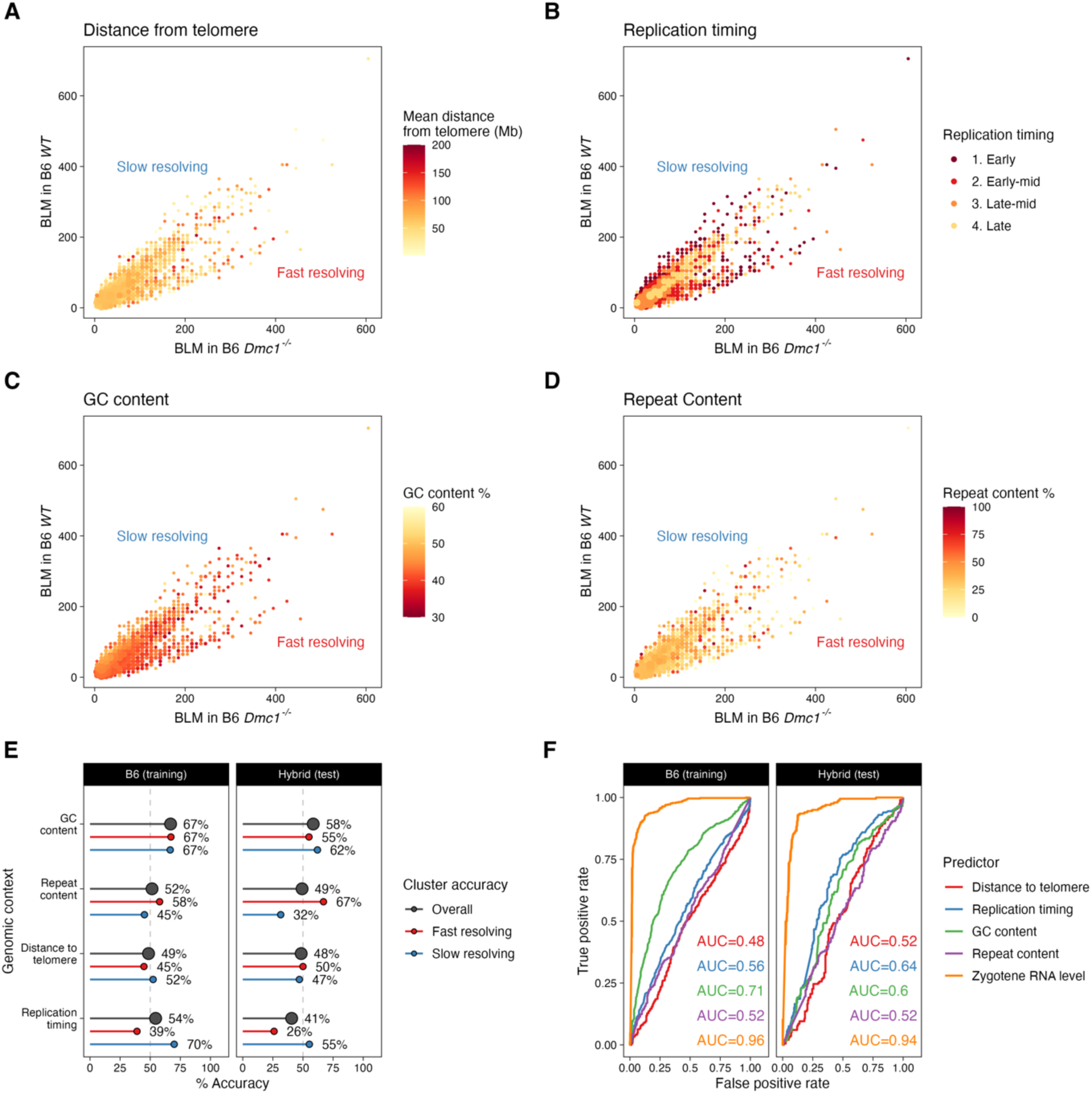
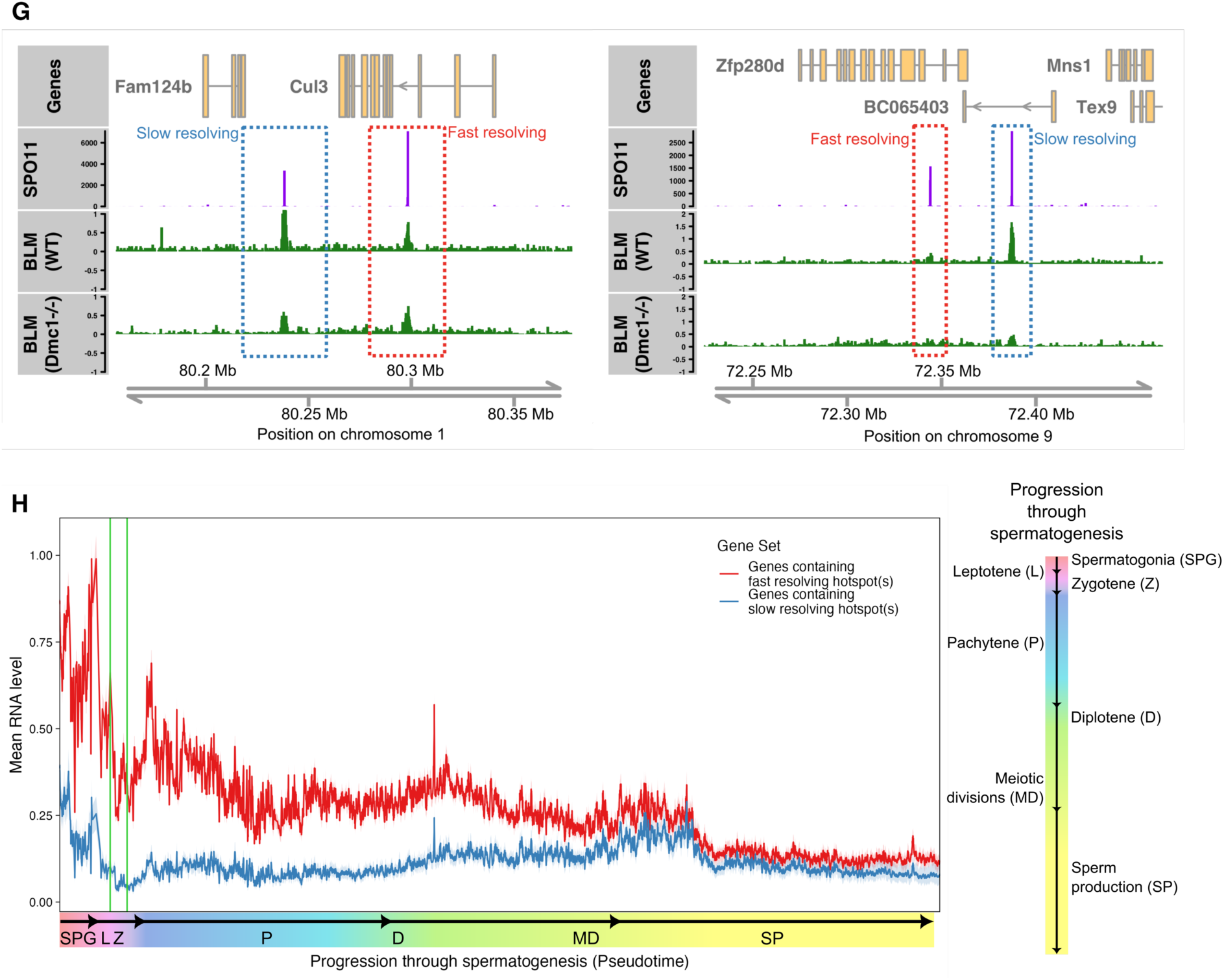
Zygotene RNA levels predict rapid strand-exchange resolution across mouse sub-species independently of genomic confounders. **(A)** Scatterplot of strand-exchange–dependent BLM signal (wild-type B6 relative to *Dmc1^-/-^*) across hotspots aggregated within cells of a 75 × 75 grid (as in Figure 3B). Each point represents one grid cell; colour indicates average distance of hotspots in that cell from the non-centromeric telomere, and point size reflects hotspot density. No systematic gradient is observed. **(B)** As in (A) but coloured by replication timing (Data from^35^), showing no consistent relationship between replication timing and strand-exchange–dependent class separation. **(C)** As in (A) but coloured by GC-content within 1 kb of hotspot centres, indicating that GC-content does not explain strand-exchange–dependent class separation. **(D)** As in (A) but coloured by the proportion of bases within 1 kb of hotspot centres annotated as repetitive elements by RepeatMasker, demonstrating that repeat density does not account for the observed separation. **(E)** Prediction accuracy for hotspots in fast- and slow-resolving classes separately (as in Figure 3A). **(F)** Receiver operator characteristic (ROC) curves for training (B6) and test (hybrid) datasets for selected predictors, demonstrating robust generalisation of Zygotene RNA level-based prediction across genetic backgrounds. **(G)** Representative fast- and slow-resolving hotspots on chromosome 1, showing SPO11-oligo and BLM ChIP-SSDS signal in WT and *Dmc1^-/-^* mice. **(H)** Average RNA level of genes harbouring fast- and slow-resolving hotspots across pseudotime in spermatogenesis (top 500 most confidently classified genes per class). Germ-cell stages, namely, spermatogonia (SPG), Leptotene (L), Zygotene (Z), Pachytene (P), Diplotene (D), Meiotic divisions (MD) & Sperm production (SP), are annotated (Data from^48^).

**Figure S4.**
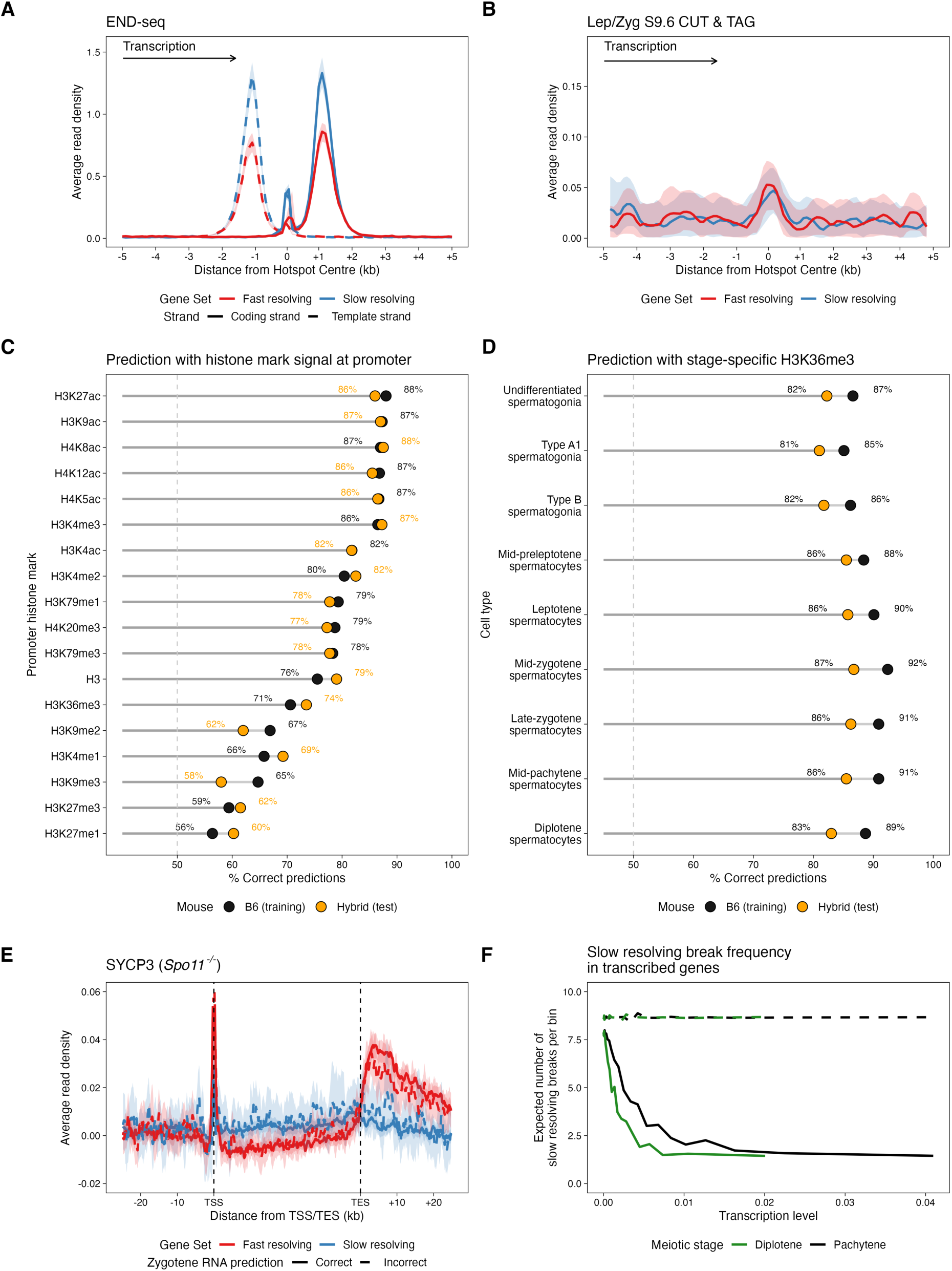
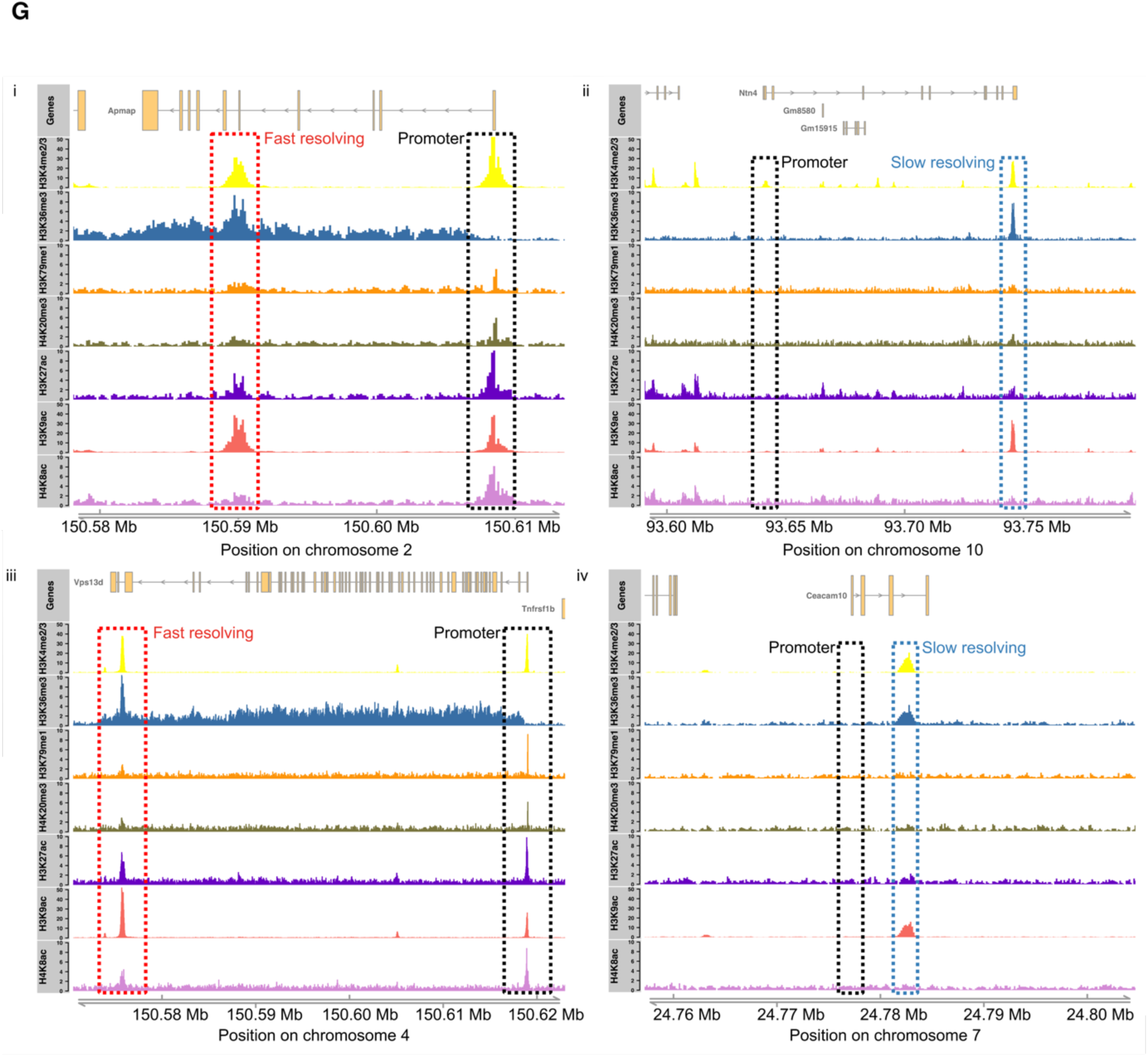
Structural and epigenetic features of prior transcription define micro-domains of rapid strand-exchange resolution. **(A)** END-seq signal in genic fast- and slow-resolving hotspots, stratified by the transcribed (‘template’, dashed) versus non-transcribed (‘coding’, solid) DNA strands (100 bp windows). END-seq detects ssDNA–dsDNA junctions generated during DSB processing and strand exchange. If rapid resolution were driven by direct transcription–recombination conflicts (e.g., due to polymerase collision), a strand-asymmetric END-seq profile would be expected. However, no systematic strand bias is observed in either fast- or slow-resolving hotspots. This suggests that the kinetic divergence does not arise from interference with ongoing transcription. **(B)** As in (A) but for average R-loop signal measured via S9.6 CUT&Tag (Data from^83^). R-loop signal is similar in both fast- and slow-resolving genes. Together with panel (A), this indicates that the repair bias is not explained by active transcription. **(C)** Prediction accuracy of transcription start site (TSS)-proximal chromatin marks for classifying genic hotspots as fast- or slow-resolving. Chromatin features at promoters are highly predictive. **(D)** Predictive accuracy of H3K36me3 measured at distinct stages of spermatogenesis (chromatin marks data from^58^). Zygotene-stage H3K36me3 provides the strongest predictive power, reinforcing stage specificity of the regulatory signal. **(E)** Metagene profiles of SYCP3 ChIP-seq signal in *Spo11^-/-^* mice, comparing genes harbouring fast- (red) and slow-resolving (blue) hotspots, stratified by classification accuracy based on zygotene RNA level. Solid lines denote genes correctly classified by RNA level; dashed lines denote misclassified genes. SYCP3 is a core component of the meiotic chromosome axis. Differences in axis association between fast- and slow-resolving genes persist in the absence of DSB formation, indicating that this structural distinction is established independently of meiotic breaks. **(F)** Expected number of slow-resolving breaks occurring within genes binned by transcription level in pachytene (black) and diplotene (green), with bins constructed to contain equal total SPO11-oligo counts. Solid lines indicate expectations derived from mid-zygotene H3K36me3-based probabilities of slow resolution, whereas dashed lines represent a (null) model in which all breaks are assumed to resolve slowly. Genes with high transcription levels in pachytene and diplotene are predicted to host very few slow-resolving breaks, consistent with rapid resolution protecting genes required for later transcriptional activity. **(G)** Representative examples of B6 fast- and slow-resolving genic hotspots shown with ChIP-seq coverage for H3K4me2/3 (yellow), H3K36me3 (blue), H3K79me1 (orange), H4K20me3 (khaki), H3K27ac (purple), H3K9ac (salmon), and H4K8ac (plum), illustrating coordinated transcription-associated chromatin signatures in fast-resolving genes.

**Figure S5.**
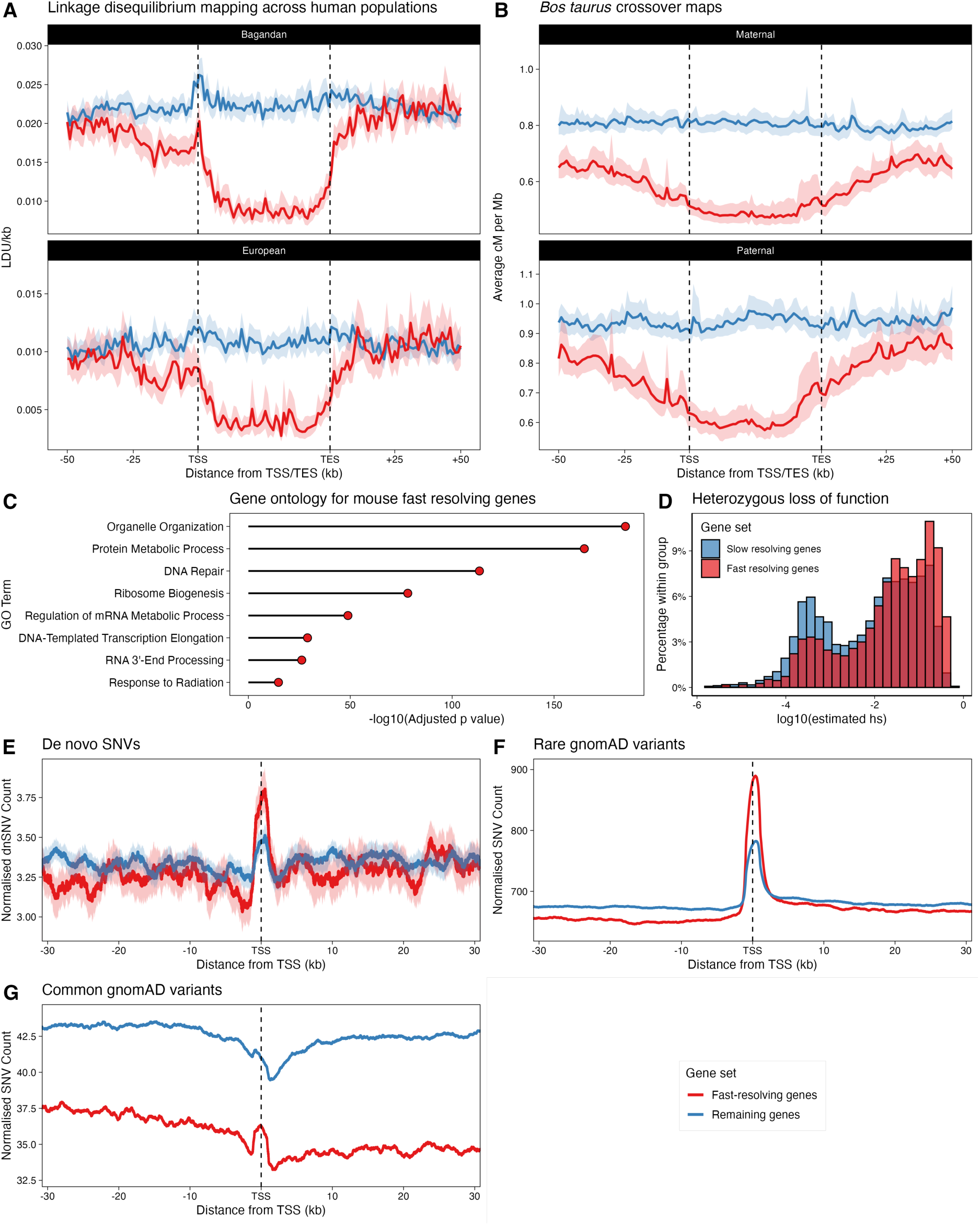
Evolutionary conservation of crossover suppression and functional constraint in fast-resolving genes. **(A)** Linkage disequilibrium unit (LDU/kb) metagene profiles across human orthologues of predicted fast-resolving genes (n=4,227; red) versus other protein-coding genes (n=13,676; blue) in an African (top) and a European (bottom) population. Gene bodies were scaled to 40kb and plotted using 1 kb bins. Reduced LDU values across fast-resolving gene bodies indicate suppressed historical recombination in both populations. **(B)** Average maternal (left) and paternal (right) crossover rates across bovine orthologues of predicted fast-resolving genes (n=3,616; red) compared with other genes (n=12,028; blue), using the same scaling and bin size as in (A) (Data from^59^). Crossover suppression within fast-resolving gene orthologues is conserved in cattle and evident in both sexes. **(C)** Gene ontology enrichment analysis for mouse genes predicted to be fast resolving, showing the most significantly enriched biological process terms. Fast-resolving genes are enriched for essential cellular and DNA repair functions. **(D)** Distribution of haploinsufficiency scores for human orthologues of predicted fast-resolving compared with other protein-coding genes (Data from^81^). Fast-resolving genes exhibit higher haploinsufficiency scores, consistent with greater functional constraint. **(E)** Density of human de novo single nucleotide variants (SNVs) per gene around transcription start sites of predicted fast-resolving genes (red) and other protein-coding genes (blue) in a multi-cohort dataset (gnomAD)^80^. Genes are oriented such that transcription proceeds from left to right (2 kb rolling smoothing window). De novo mutation rates are broadly comparable between these gene sets. **(F)** As in (E) but for rare SNVs (minor allele frequency < 10^-4^) in gnomAD. Fast-resolving genes exhibit broadly comparable rare variant density, consistent with similar mutation rates. **(G)** As in (E) but for common SNVs (minor allele frequency > 10^-2^) in gnomAD. Strong reduction in common variant density in fast-resolving genes is consistent with crossover suppression and increased selective constraint.

**Figure S6.**
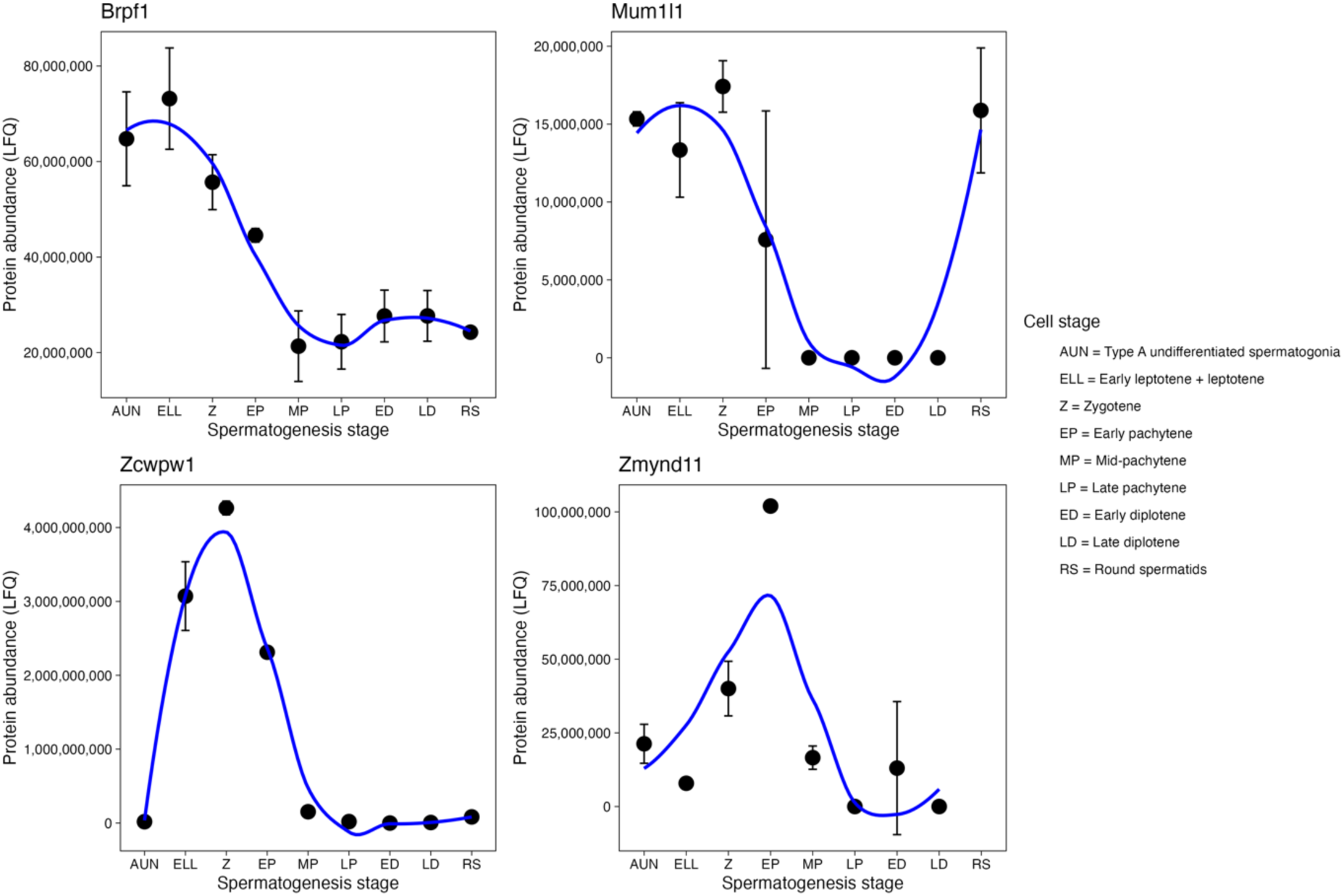
Stage-specific abundance of candidate H3K36me3-binding proteins across spermatogenesis. Stage-specific abundance of BRPF1 (top left), MUM1L1/PWWP3B (top right), ZCWPW1 (bottom left) and ZMYND11 (bottom right), measured by label-free quantification across spermatogenic stages. Data from^85^. These proteins contain PWWP or H3K36me3-recognition domains and are expressed during meiotic prophase, supporting the availability of candidate chromatin readers that could interpret H3K36me3-enriched domains during strand-exchange resolution. See Supplementary Text for further discussion.

**Table S1.** Key metrics for BLM maps generated through ChIP-SSDS in adult mouse testes. The two BLM replicates in the hybrid showed high signal concordance over 24,586 hotspots in the hybrid (Pearson’s r=0.98). Data from the highlighted replicates are reported in the paper.

|  | **BLM in B6** | **BLM in B6**<br>***Dmc1*<sup>-/-</sup>** | **BLM in**<br>**hybrid**<br>**(replicate**<br>**1)** | BLM in<br>hybrid<br>(replicate 2) | **BLM in**<br>**hybrid**<br>***Dmc1*<sup>-/-</sup>** |
| --- | --- | --- | --- | --- | --- |
| Number of reads | 344,671,542 | 380,495,160 | 235,761,650 | 199,944,576 | 282,508,844 |
| Percentage of reads mapped to reference (mm10) | 91% | 86% | 87% | 66% | 74% |
| Number of ssDNA (Type 1) reads identified | 81,389,803 | 71,730,401 | 49,076,309 | 23,149,391 | 41,714,934 |
| Number of ssDNA (Type 1) reads inferred to be unique | 10,145,496 | 24,342,853 | 6,602,931 | 5,255,956 | 1,603,816 |

**Table S2.**
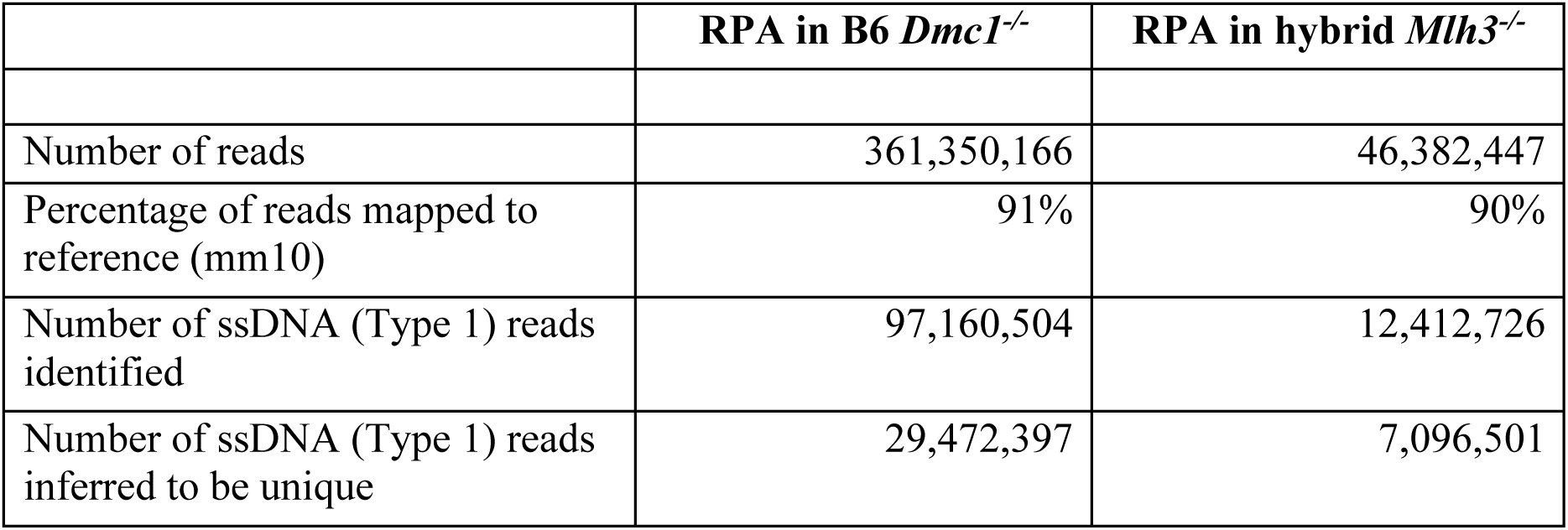
Key metrics for RPA maps generated following ChIP-SSDS in adult mouse testes.

**Table S3.** Key metrics for HFM1 maps generated following ChIP-SSDS in adult mouse testes. Data from the highlighted replicates are reported in the paper.

|  | **HFM1 in B6** | HFM1 in B6 | **HFM1 in hybrid** | HFM1 in hybrid |
| --- | --- | --- | --- | --- |
|  | (replicate 1) | (pool of 3 replicates) | (replicate 1) | (pool of 5 replicates) |
| Number of reads | 314,774,102 | 199,343,106 | 394,793,694 | 216,399,698 |
| Percentage of reads mapped to reference (mm10) | 90% | 87% | 86% | 88% |
| Number of ssDNA (Type 1) reads identified | 86,445,411 | 46,378,833 | 101,543,068 | 49,703,498 |
| Number of ssDNA (Type 1) reads inferred to be unique | 47,802,507 | 20,381,779 | 66,011,493 | 23,943,308 |

**Table S4.**
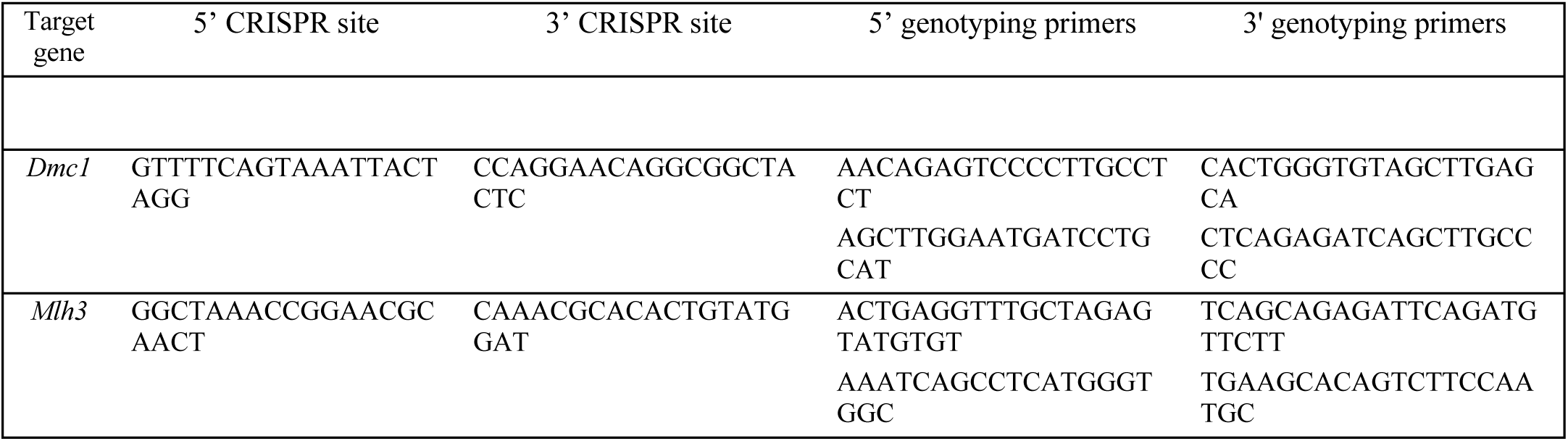
CRISPR target site and genotyping information for the creation and validation of *Dmc1* and *Mlh3* mouse knockouts.

**Table S5.** Fertility parameters and immunohistochemistry (IHC) assessment of putative *Dmc1* and *Mlh3* knockout mice. Spread spermatocytes were stained for DMC1 (in *Dmc1^-/^*^-^) and MLH1 (in *Mlh3^-/^*^-^) and assessed for the presence of foci in leptotene and pachytene cells, respectively. No sperm were present and no cells with DMC1 (in *Dmc1^-/^*^-^) or MLH1 (in *Mlh3^-/-^*) foci were detected, consistent with complete ablation of DMC1 and MLH3 respectively.

| Mouse ID | Gene KO | Genetic background | Cells with DMC1 foci<br>/<br>Number assessed | Cells with MLH1 foci<br>/<br>Number assessed | Sperm count | ChIP-SSDS |
| --- | --- | --- | --- | --- | --- | --- |
| TGMS4.1a | *Mlh3* | B6CASTF1 | - | 0/102 | 0 | RPA |
| TGMS8.1a | *Mlh3* | B6CASTF1 | - | 0/103 | 0 |  |
| TGMZ9.1c | *Dmc1* | C57BL/6J | 0/83 | - | 0 | RPA |
| TGMZ17.1<br>e | *Dmc1* | B6CASTF1 | Described in<sup>43</sup> |  |  | BLM |
| TGMZ10.1<br>a | *Dmc1* | B6CASTF1 | Described in<sup>37</sup> |  |  | BLM |

### Supplementary Text

#### BLM Clusters Reflect Differences in the Lifespan of Strand-Exchange Intermediates

Segregation of wild-type hotspots into clusters with systematically high and low BLM signal could, in principle, arise from several distinct phenomena. Differences in BLM occupancy could reflect:

1. variation in BLM loading onto recombination intermediates;
2. differences in the lifespan (persistence) of strand-exchange intermediates; or
3. artefactual effects arising from read mappability, antibody accessibility, or antibody affinity.

To distinguish between these possibilities, we carried out a series of analyses designed to test the predictions of each hypothesis. As a first step, we quantified the strand-exchange component as follows.

### Quantification of strand-exchange component

Because ChIP-SSDS read counts depend on experimental factors such as sequencing depth and the number of cells processed, absolute coverage values are not directly comparable between wild-type and *Dmc1*^⁻/⁻^ assays. In contrast, relative differences between hotspots *within* each experiment are preserved. For this reason, we used the ratio of wild-type to *Dmc1*^⁻/⁻^ BLM signal at each hotspot to quantify the strand-exchange–dependent component. This approach normalises global scaling differences between experiments while retaining biologically meaningful hotspot-to-hotspot variation attributable to strand exchange.

To ensure that the hotspots were directly comparable for break frequency, we generated sets of hotspots matched for SPO11-oligo density (Figure 1F, STAR Methods). Matching break frequency allows us to test which subsequent step in the recombination process drives the difference in BLM occupancy.

### 1. Protein-independent measurements

We examined published data from END-seq, which measures ssDNA-dsDNA junctions directly without antibodies, in B6 and mutant mouse testes as follows:

**1.1. END-seq ssDNA-dsDNA junctions show similar resection lengths in the hotspot clusters.** If the clusters reflected differences in resected ssDNA production, the shape and width of END-seq would be different in the flanking (non-central region) between the clusters. However, the footprint shape and width were indistinguishable between the hotspot classes in both the wild-type and strand-exchange mutants (Figure S1J-L). Therefore, there is no evidence that slow-resolving hotspots undergo greater resection or produce more ssDNA before strand invasion.
**1.2. END-seq in wild type shows greater D-loop signal at slow-resolving hotspots.** The central signal in END-seq reflects strand-exchange intermediates^42^. In wild-type testes, END-seq revealed a larger central ssDNA signal at slow-resolving hotspots (Figure S1J). Since the number of breaks is similar in the two groups, this finding is consistent with a longer persistence of D-loop structures in slow-resolving hotspots. END-seq also showed greater ssDNA signal in flanking regions at these hotspots, consistent with ssDNA in strand-exchange intermediates^43^.
**1.3. END-seq differences collapse completely in strand-exchange mutants.** In *Dmc1*^⁻/⁻^ and *Hop2*^⁻/⁻^, testes, each of which fails to form D-loops, END-seq showed no difference between the hotspot classes (Figure S1K-L). When strand invasion fails, ssDNA arises only from resection, and the clusters become indistinguishable.

Thus, protein-independent measures show the same separation as BLM only when D-loops form, and this separation disappears when strand exchange is abolished. These results rule out differential BLM loading and technical artefacts, because the cluster separation is reproduced by a completely antibody-independent assay and vanishes when D-loops cannot form.

### 2. Resection length does not differ between clusters

If the clusters reflected differences in upstream ssDNA production, then resection lengths or ssDNA footprints should differ systematically. We have already considered END-seq ssDNA-dsDNA junction distribution above. Here we examine three further measures of resection: footprint widths of BLM, RPA, and DMC1.

In all cases, the footprint shape and width were indistinguishable between the hotspot classes (Figure 1G-H, Figure S1H, M-N). As with END-seq, there is no evidence that slow-resolving hotspots undergo greater resection or produce more ssDNA before strand invasion. Therefore, cluster separation cannot be attributed to differences in the amount or distribution of resected ssDNA.

### 3. Homology search efficiency is equivalent between clusters

We next assessed homology search using DMC1 occupancy. After matching for number of breaks, DMC1 level is informative about the duration of homology search (homologue-engagement time)^36,43^. Homologue-engagement was indistinguishable between fast- and slow-resolving hotspots (Figure 1G). This rules out systematic differences in homology search efficiency, availability of donor templates, or frequency of strand invasion attempts, none of which can therefore explain the BLM cluster separation.

### 4. RPA differences collapse in strand-exchange mutants

RPA, like BLM, binds both resected ssDNA and strand-exchange intermediates. In wild type, RPA levels differ between the two hotspot classes (Figure S1M). However, in *Dmc1*^⁻/⁻^ testes, RPA differences disappear completely (Figure S1N). These results are again consistent with clusters being different due to differences in the persistence of strand-exchange intermediates.

### Integrating all the evidence: cluster separation reflects lifespan of strand-exchange intermediates

Across all assays: BLM ChIP-SSDS, RPA ChIP-SSDS, END-seq, DMC1 ChIP-SSDS, and analyses of *Dmc1*^⁻/⁻^ and *Hop2*^⁻/⁻^ mutants, a single coherent explanation accounts for the observations:

- Differences between clusters are present only in wild type, when D-loops form and persist.
- Differences disappear in each genotype where D-loop formation fails.
- Differences are not explained by break number, resection extent, or homology search efficiency.
- Differences persist after controlling for mappability, antibody affinity, and antibody accessibility.
- Differences are replicated qualitatively and quantitatively across independent assays and proteins.

Taken together, these analyses eliminate hypotheses based on BLM loading or technical artefacts such as read mappability, antibody affinity, or accessibility. They establish that BLM occupancy reports persistence of strand-exchange intermediates, and the clustering therefore reflects a biologically meaningful kinetic distinction in meiotic repair.

## Statistical Framework for Unsupervised Clustering of Hotspots Into Two Classes

Repair of meiotic DSBs generates both early single-stranded DNA intermedi-ates from resection and later intermediates from strand-exchange (D-loops). In *Dmc*1*^-/-^* mice, strand invasion fails and BLM SSDS signal reflects only early intermediates. In wild type, the same assays capture both early and post-strand-exchange intermediates. Comparing wild-type to *Dmc*1*^-/-^* signals therefore allows inference of the relative persistence of strand-exchange intermediates at each hotspot.

Exploratory analysis of wild-type versus *Dmc*1*^-/-^* BLM signal revealed two visually distinct clusters of hotspots. We therefore sought to determine, in an unsupervised and data-driven manner, whether these clusters represent genuine classes with systematically different properties, and to compute for each hotspot the posterior probability of belonging to either class. No hotspot was pre-assigned to a cluster; the model learns class membership solely from the data, given the assumption of two latent groups.

Since we have multiple experimental assays informative about the persistence of strand-exchange intermediates, we sought to leverage them jointly. Specifically, for each hotspot, we have data from BLM and RPA ChIP-SSDS measuring their occupancy, which reflects both loading and persistence, at that location in both wild-type and *Dmc*1*^-/-^* mice. We refer to this as ’activity’ henceforth.

Formally, we assign one experiment as measuring the reference level of activity for the location, with the other experiments measuring different aspects of activity. Inspection of the data suggests that there are two classes of location, one with a high activity relative to the reference and one with low activity relative to the reference. For each assay and class, the experimental count data increases approximately linearly with the reference level. Whether a location is in the high or low activity class is assumed to be the same across all experiments, with each assay providing as noisy estimate of the activity. Our task is to infer the class of each location by combining the results of all the experimental assays. We take a Bayesian hierarchical approach which combines a mixture model for class membership with Poisson regression as follows.

Define *N* as the total number of hotspots and *M* as the number of experimental assays measuring activity. The reference data for each location is *x_i_* where i 2 *{*1, ..N *}*; and the experiment count data for each assay is *y_i,j_* where *j* ∈ *{*1, ..M *}*. Define *Z_i_* as an indicator variable of class memberships, with *Z_i_* =0 being the low activity class and *Z_i_* =1 the high activity class. The experimental count data (*Y_i,j_*) are modelled as independent Poisson variables

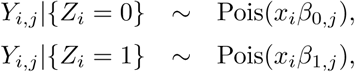

where β_0*,j*_ and β_1*,j*_ are regression parameters which are estimated. The experi-mental counts for each assay are assumed to be independent, so

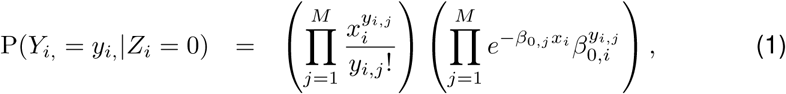

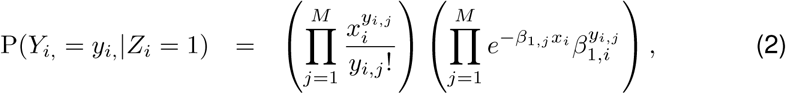

where *Y_i,_* = *{Y_i,_*_1_, .., *Y_i,M_ }*. Note that the first product is the same for both classes and independent of the regression parameters, therefore can be dropped when sampling the posterior distribution using MCMC. The law of total probability gives

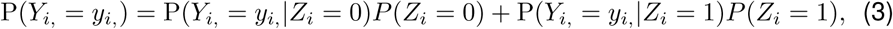

where *P*(*Z_i_* = 0) is the prior probability that the i^th^ location is in the low activity state. We now assume that this prior probability is the independent of location, so

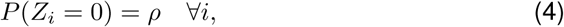

where *ρ* is the prior probability which is estimated across all data (*i.e.* it is a hierarchical Bayesian model}. Combining (1), (3) and (4) gives

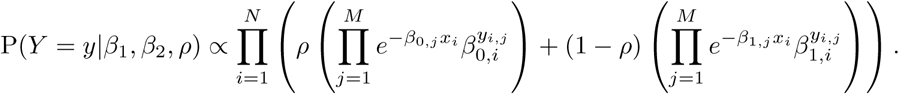

Finally, we use range priors on *β*_1_, *β*_2_ and *ρ*, and Bayes theorem to get the posterior distribution

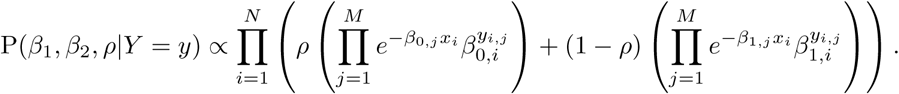

which we sample from using MCMC. The probability of class membership of a location given the count data and model parameters is

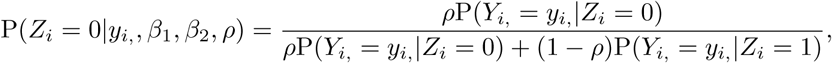

which can then be integrated over the posterior distribution of {*β*_1_, *β*_2_, *ρ*} sampled using Markov Chain Monte Carlo (MCMC).

The model was implemented in Stan and run using the default settings for B6 and hybrid hotspots separately. The Poisson count model led to an under-estimation of the experimental error for counts at hotspots with high reference levels, therefore we performed a variance-stabilising transform on the counts before applying the model (power 0.6 for B6 and 0.7 for hybrid). Hotspots with reference levels below a threshold (weak hotspots) were excluded due to insufficient information for clustering (cutoffs were 300 for B6 and 280 for hybrid).

### Constraints and potential molecular mechanisms of transcription-linked repair partitioning

The analyses presented in this study place strong constraints on the molecular mechanisms that could underlie the observed partitioning of meiotic DSB repair outcomes. Any viable mechanism must operate at the level of gene-intrinsic properties rather than at individual hotspots, generalise across genomes with distinct break landscapes, and act at or before the stage at which repair pathways diverge. Moreover, it must be compatible with rapid turnover of strand-exchange intermediates at fast-resolving loci without impairing homology search or early pairing.

Below, we outline classes of molecular mechanisms that are consistent with these constraints. These possibilities are not mutually exclusive and there may be complementary routes by which transcriptional memory could influence meiotic repair dynamics.

#### Chromatin-linked mechanisms

One potential mechanism involves chromatin states associated with prior transcription, particularly elongation-associated histone modifications such as H3K36me3. This mark is deposited co-transcriptionally, persists after transcriptional repression in early meiotic prophase, and strongly predicts fast resolution of strand-exchange intermediates in our data. A simple interpretation is that H3K36me3-rich chromatin may alter the local environment of recombination intermediates, for example by modulating stability of repair factors. To explore this possibility, we examined the proteomic time courses^85^ of known readers of H3K36me3 and/or transcriptionally-primed chromatin across meiotic progression:

We prioritised genes with high proteomic measurements in zygotene. From the initial set of candidate proteins, proteomic signatures favour BRPF1, MUM1L1(PWWP3B), ZCWPW1, and ZMYND11 (Figure S6).

#### Structural and axis-associated mechanisms

An alternative, or complementary, class of mechanisms involves higher-order chromosome organisation. Meiotic chromosomes are arranged into loop-axis structures, and regulatory elements such as transcription start sites (TSS) are known to associate with the chromosome axis^55,86^. RNA polymerase II, paused near the TSS of thousands of genes during early prophase^51^, interacts with SYCP3^86^, providing a direct link between loop-axis organisation and transcription. Our observation that fast-resolving genes exhibit axis association at both transcription start and termination sites suggests that entire gene bodies may form constrained loop domains during early prophase.

Such an arrangement could influence repair dynamics by limiting the extension or stabilisation of strand-exchange intermediates, or by restricting access of factors required for crossover designation. Recent studies in budding yeast demonstrate that key components of the synaptonemal complex protect crossover-designated recombination intermediates from non-crossover processing, highlighting how higher-order chromosome architecture can regulate repair pathway choice^56,57^. Importantly, this interpretation does not require chromatin state to act independently of structure; rather, chromatin and architecture may jointly encode transcriptional memory through stable gene-scale organisation.

Consistent with this view, several proteins implicated in stabilising recombination intermediates interact with axis components. For example, HFM1 and RNF212, are associated with intermediate stabilisation and crossover designation and identified in human GWAS as modifiers of the chromatin context of crossovers^14^. These proteins interact with components of the synaptonemal complex^31,87,88^, which provides a plausible molecular bridge between gene-level organisation and the differential persistence of recombination intermediates.

#### Integration of chromatin and structural features

The data presented here do not distinguish whether chromatin state or chromosome architecture is primary. Indeed, it is possible that transcriptional memory is implemented through an interplay of both. Elongation-associated chromatin marks may promote or stabilise specific modes of axis engagement, which in turn may shape the spatial constraints experienced by recombination intermediates. In this view, transcriptional history is encoded not in a single molecular mark or factor, but in a composite gene-intrinsic state that persists through early meiotic prophase and biases repair dynamics once breaks occur.

Future experiments that selectively alter chromatin state, axis association, or their coupling, will be required to dissect how transcriptional memory is implemented at the molecular level. The framework outlined here provides a set of experimentally testable constraints within which such mechanisms would operate.

## References

1. Hunter, N. (2015). Meiotic Recombination: The Essence of Heredity. Cold Spring Harb Perspect Biol 7. 10.1101/cshperspect.a016618.

2. Coop, G., and Przeworski, M. (2007). An evolutionary view of human recombination. Nat Rev Genet 8, 23–34. 10.1038/nrg1947.

3. Petronczki, M., Siomos, M.F., and Nasmyth, K. (2003). Un Ménage à Quatre: The Molecular Biology of Chromosome Segregation in Meiosis. Cell 112, 423–440. 10.1016/S0092-8674(03)00083-7.

4. Baudat, F., Imai, Y., and de Massy, B. (2013). Meiotic recombination in mammals: localization and regulation. Nat Rev Genet 14, 794–806. 10.1038/nrg3573.

5. Baudat, F., and de Massy, B. (2007). Regulating double-stranded DNA break repair towards crossover or non-crossover during mammalian meiosis. Chromosome Res 15, 565–577. 10.1007/s10577-007-1140-3.

6. Awwad, S.W., Abu-Zhayia, E.R., Guttmann-Raviv, N., and Ayoub, N. (2017). NELF-E is recruited to DNA double-strand break sites to promote transcriptional repression and repair. EMBO Rep 18, 745–764. 10.15252/embr.201643191.

7. Djeghmoum, Y., and Piazza, A. (2025). Donor transcription suppresses D-loops in cis and promotes genome stability. Embo j 44, 5595–5617. 10.1038/s44318-025-00541-x.

8. Kruhlak, M., Crouch, E.E., Orlov, M., Montaño, C., Gorski, S.A., Nussenzweig, A., Misteli, T., Phair, R.D., and Casellas, R. (2007). The ATM repair pathway inhibits RNA polymerase I transcription in response to chromosome breaks. Nature 447, 730–734. 10.1038/nature05842.

9. Shanbhag, N.M., Rafalska-Metcalf, I.U., Balane-Bolivar, C., Janicki, S.M., and Greenberg, R.A. (2010). ATM-dependent chromatin changes silence transcription in cis to DNA double-strand breaks. Cell 141, 970–981. 10.1016/j.cell.2010.04.038.

10. Hinch, R., Donnelly, P., and Hinch, A.G. (2023). Meiotic DNA breaks drive multifaceted mutagenesis in the human germ line. Science 382, eadh2531. 10.1126/science.adh2531.

11. Davies, B., Hatton, E., Altemose, N., Hussin, J.G., Pratto, F., Zhang, G., Hinch, A.G., Moralli, D., Biggs, D., Diaz, R., et al. (2016). Re-engineering the zinc fingers of PRDM9 reverses hybrid sterility in mice. Nature 530, 171–176. 10.1038/nature16931.

12. McVicker, G., and Green, P. (2010). Genomic signatures of germline gene expression. Genome Res 20, 1503–1511. 10.1101/gr.106666.110.

13. Palsson, G., Hardarson, M.T., Jonsson, H., Steinthorsdottir, V., Stefansson, O.A., Eggertsson, H.P., Gudjonsson, S.A., Olason, P.I., Gylfason, A., Masson, G., et al. (2025). Complete human recombination maps. Nature 639, 700–707. 10.1038/s41586-024-08450-5.

14. Halldorsson, B.V., Palsson, G., Stefansson, O.A., Jonsson, H., Hardarson, M.T., Eggertsson, H.P., Gunnarsson, B., Oddsson, A., Halldorsson, G.H., Zink, F., et al. (2019). Characterizing mutagenic effects of recombination through a sequence-level genetic map. Science 363. 10.1126/science.aau1043.

15. Jin, X., Fudenberg, G., and Pollard, K.S. (2021). Genome-wide variability in recombination activity is associated with meiotic chromatin organization. Genome Res 31, 1561–1572. 10.1101/gr.275358.121.

16. Soumillon, M., Necsulea, A., Weier, M., Brawand, D., Zhang, X., Gu, H., Barthes, P., Kokkinaki, M., Nef, S., Gnirke, A., et al. (2013). Cellular source and mechanisms of high transcriptome complexity in the mammalian testis. Cell Rep 3, 2179–2190. 10.1016/j.celrep.2013.05.031.

17. Geisinger, A., Rodriguez-Casuriaga, R., and Benavente, R. (2021). Transcriptomics of Meiosis in the Male Mouse. Front Cell Dev Biol 9, 626020. 10.3389/fcell.2021.626020.

18. Shima, J.E., McLean, D.J., McCarrey, J.R., and Griswold, M.D. (2004). The murine testicular transcriptome: characterizing gene expression in the testis during the progression of spermatogenesis. Biol Reprod 71, 319–330. 10.1095/biolreprod.103.026880.

19. Xia, B., Yan, Y., Baron, M., Wagner, F., Barkley, D., Chiodin, M., Kim, S.Y., Keefe, D.L., Alukal, J.P., Boeke, J.D., and Yanai, I. (2020). Widespread Transcriptional Scanning in the Testis Modulates Gene Evolution Rates. Cell 180, 248–262.e221. 10.1016/j.cell.2019.12.015.

20. Hatkevich, T., and Sekelsky, J. (2017). Bloom syndrome helicase in meiosis: Pro-crossover functions of an anti-crossover protein. Bioessays 39. 10.1002/bies.201700073.

21. Holloway, J.K., Morelli, M.A., Borst, P.L., and Cohen, P.E. (2010). Mammalian BLM helicase is critical for integrating multiple pathways of meiotic recombination. J Cell Biol 188, 779–789. 10.1083/jcb.200909048.

22. Powers, N.R., Parvanov, E.D., Baker, C.L., Walker, M., Petkov, P.M., and Paigen, K. (2016). The Meiotic Recombination Activator PRDM9 Trimethylates Both H3K36 and H3K4 at Recombination Hotspots In Vivo. PLoS Genet 12, e1006146. 10.1371/journal.pgen.1006146.

23. Eram, M.S., Bustos, S.P., Lima-Fernandes, E., Siarheyeva, A., Senisterra, G., Hajian, T., Chau, I., Duan, S., Wu, H., Dombrovski, L., et al. (2014). Trimethylation of histone H3 lysine 36 by human methyltransferase PRDM9 protein. J Biol Chem 289, 12177–12188. 10.1074/jbc.M113.523183.

24. Baudat, F., Buard, J., Grey, C., Fledel-Alon, A., Ober, C., Przeworski, M., Coop, G., and de Massy, B. (2010). PRDM9 is a major determinant of meiotic recombination hotspots in humans and mice. Science 327, 836–840. 10.1126/science.1183439.

25. Lange, J., Yamada, S., Tischfield, S.E., Pan, J., Kim, S., Zhu, X., Socci, N.D., Jasin, M., and Keeney, S. (2016). The Landscape of Mouse Meiotic Double-Strand Break Formation, Processing, and Repair. Cell 167, 695–708.e616. 10.1016/j.cell.2016.09.035.

26. Symington, L.S. (2016). Mechanism and regulation of DNA end resection in eukaryotes. Crit Rev Biochem Mol Biol 51, 195–212. 10.3109/10409238.2016.1172552.

27. Crickard, J.B., and Greene, E.C. (2018). Biochemical attributes of mitotic and meiotic presynaptic complexes. DNA Repair (Amst) 71, 148–157. 10.1016/j.dnarep.2018.08.018.

28. Hinch, A.G., Becker, P.W., Li, T., Moralli, D., Zhang, G., Bycroft, C., Green, C., Keeney, S., Shi, Q., Davies, B., and Donnelly, P. (2020). The Configuration of RPA, RAD51, and DMC1 Binding in Meiosis Reveals the Nature of Critical Recombination Intermediates. Mol Cell 79, 689-701 e610. 10.1016/j.molcel.2020.06.015.

29. Shorrocks, A.-M.K., Jones, S.E., Tsukada, K., Morrow, C.A., Belblidia, Z., Shen, J., Vendrell, I., Fischer, R., Kessler, B.M., and Blackford, A.N. (2021). The Bloom syndrome complex senses RPA-coated single-stranded DNA to restart stalled replication forks. Nature Communications 12, 585. 10.1038/s41467-020-20818-5.

30. Altmannova, V., Firlej, M., Müller, F., Janning, P., Rauleder, R., Rousova, D., Schäffler, A., Bange, T., and Weir, J.R. (2023). Biochemical characterisation of Mer3 helicase interactions and the protection of meiotic recombination intermediates. Nucleic Acids Res 51, 4363–4384. 10.1093/nar/gkad175.

31. Guiraldelli, M.F., Eyster, C., Wilkerson, J.L., Dresser, M.E., and Pezza, R.J. (2013). Mouse HFM1/Mer3 Is Required for Crossover Formation and Complete Synapsis of Homologous Chromosomes during Meiosis. PLOS Genetics 9, e1003383. 10.1371/journal.pgen.1003383.

32. Mazina, O.M., Mazin, A.V., Nakagawa, T., Kolodner, R.D., and Kowalczykowski, S.C. (2004). Saccharomyces cerevisiae Mer3 helicase stimulates 3’-5’ heteroduplex extension by Rad51; implications for crossover control in meiotic recombination. Cell 117, 47–56. 10.1016/s0092-8674(04)00294-6.

33. Gray, S., and Cohen, P.E. (2016). Control of Meiotic Crossovers: From Double-Strand Break Formation to Designation. Annu Rev Genet 50, 175–210. 10.1146/annurev-genet-120215-035111.

34. de Boer, E., Jasin, M., and Keeney, S. (2015). Local and sex-specific biases in crossover vs. noncrossover outcomes at meiotic recombination hot spots in mice. Genes Dev 29, 1721–1733. 10.1101/gad.265561.115.

35. Pratto, F., Brick, K., Cheng, G., Lam, K.G., Cloutier, J.M., Dahiya, D., Wellard, S.R., Jordan, P.W., and Camerini-Otero, R.D. (2021). Meiotic recombination mirrors patterns of germline replication in mice and humans. Cell 184, 4251–4267.e4220. 10.1016/j.cell.2021.06.025.

36. Hinch, A.G., Zhang, G., Becker, P.W., Moralli, D., Hinch, R., Davies, B., Bowden, R., and Donnelly, P. (2019). Factors influencing meiotic recombination revealed by whole-genome sequencing of single sperm. Science 363. 10.1126/science.aau8861.

37. Li, R., Bitoun, E., Altemose, N., Davies, R.W., Davies, B., and Myers, S.R. (2019). A high-resolution map of non-crossover events reveals impacts of genetic diversity on mammalian meiotic recombination. Nature Communications 10, 3900. 10.1038/s41467-019-11675-y.

38. Serrentino, M.-E., Chaplais, E., Sommermeyer, V., and Borde, V. (2013). Differential Association of the Conserved SUMO Ligase Zip3 with Meiotic Double-Strand Break Sites Reveals Regional Variations in the Outcome of Meiotic Recombination. PLOS Genetics 9, e1003416. 10.1371/journal.pgen.1003416.

39. Khil, P.P., Smagulova, F., Brick, K.M., Camerini-Otero, R.D., and Petukhova, G.V. (2012). Sensitive mapping of recombination hotspots using sequencing-based detection of ssDNA. Genome Research 22, 957–965. 10.1101/gr.130583.111.

40. Pittman, D.L., Cobb, J., Schimenti, K.J., Wilson, L.A., Cooper, D.M., Brignull, E., Handel, M.A., and Schimenti, J.C. (1998). Meiotic prophase arrest with failure of chromosome synapsis in mice deficient for Dmc1, a germline-specific RecA homolog. Mol Cell 1, 697–705. 10.1016/s1097-2765(00)80069-6.

41. Yoshida, K., Kondoh, G., Matsuda, Y., Habu, T., Nishimune, Y., and Morita, T. (1998). The mouse RecA-like gene Dmc1 is required for homologous chromosome synapsis during meiosis. Mol Cell 1, 707–718. 10.1016/s1097-2765(00)80070-2.

42. Paiano, J., Wu, W., Yamada, S., Sciascia, N., Callen, E., Paola Cotrim, A., Deshpande, R.A., Maman, Y., Day, A., Paull, T.T., and Nussenzweig, A. (2020). ATM and PRDM9 regulate SPO11-bound recombination intermediates during meiosis. Nature Communications 11, 857. 10.1038/s41467-020-14654-w.

43. Davies, B., Zhang, G., Moralli, D., Alghadban, S., Biggs, D., Preece, C., Donnelly, P., and Hinch, A.G. (2023). Characterization of meiotic recombination intermediates through gene knockouts in founder hybrid mice. Genome Res 33, 2018–2027. 10.1101/gr.278024.123.

44. Allers, T., and Lichten, M. (2001). Differential timing and control of noncrossover and crossover recombination during meiosis. Cell 106, 47–57. 10.1016/s0092-8674(01)00416-0.

45. Börner, G.V., Kleckner, N., and Hunter, N. (2004). Crossover/noncrossover differentiation, synaptonemal complex formation, and regulatory surveillance at the leptotene/zygotene transition of meiosis. Cell 117, 29–45. 10.1016/s0092-8674(04)00292-2.

46. Cole, F., Kauppi, L., Lange, J., Roig, I., Wang, R., Keeney, S., and Jasin, M. (2012). Homeostatic control of recombination is implemented progressively in mouse meiosis. Nat Cell Biol 14, 424–430. 10.1038/ncb2451.

47. Premkumar, T., Paniker, L., Kang, R., Biot, M., Humphrey, E., Destain, H., Ferranti, I., Okulate, I., Nguyen, H., Kilaru, V., et al. (2023). Genetic dissection of crossover mutants defines discrete intermediates in mouse meiosis. Molecular Cell 83, 2941–2958.e2947. 10.1016/j.molcel.2023.07.022.

48. Jung, M., Wells, D., Rusch, J., Ahmad, S., Marchini, J., Myers, S.R., and Conrad, D.F. (2019). Unified single-cell analysis of testis gene regulation and pathology in five mouse strains. Elife 8. 10.7554/eLife.43966.

49. Goutelle, S., Maurin, M., Rougier, F., Barbaut, X., Bourguignon, L., Ducher, M., and Maire, P. (2008). The Hill equation: a review of its capabilities in pharmacological modelling. Fundamental & Clinical Pharmacology 22, 633–648. 10.1111/j.1472-8206.2008.00633.x.

50. Bellutti, L., Chan Sock Peng, E., Cluzet, V., Guerquin, M.J., Rolland, A., Messiaen, S., Llano, E., Dereli, I., Martini, E., Toth, A., et al. (2025). Genome-wide transcriptional silencing and mRNA stabilization allow the coordinated expression of the meiotic program in mice. Nucleic Acids Res 53. 10.1093/nar/gkaf146.

51. Alexander, A.K., Rice, E.J., Lujic, J., Simon, L.E., Tanis, S., Barshad, G., Zhu, L., Lama, J., Cohen, P.E., and Danko, C.G. (2023). A-MYB and BRDT-dependent RNA Polymerase II pause release orchestrates transcriptional regulation in mammalian meiosis. Nat Commun 14, 1753. 10.1038/s41467-023-37408-w.

52. Kaye, E.G., Basavaraju, K., Nelson, G.M., Zomer, H.D., Roy, D., Joseph, II, Rajabi-Toustani, R., Qiao, H., Adelman, K., and Reddi, P.P. (2024). RNA polymerase II pausing is essential during spermatogenesis for appropriate gene expression and completion of meiosis. Nat Commun 15, 848. 10.1038/s41467-024-45177-3.

53. Page, J., de la Fuente, R., Manterola, M., Parra, M.T., Viera, A., Berríos, S., Fernández-Donoso, R., and Rufas, J.S. (2012). Inactivation or non-reactivation: what accounts better for the silence of sex chromosomes during mammalian male meiosis? Chromosoma 121, 307–326. 10.1007/s00412-012-0364-y.

54. Ajit, K., and Gullerova, M. (2024). From silence to symphony: transcriptional repression and recovery in response to DNA damage. Transcription 15, 161–175. 10.1080/21541264.2024.2406717.

55. Biot, M., Toth, A., Brun, C., Guichard, L., de Massy, B., and Grey, C. (2024). Principles of chromosome organization for meiotic recombination. Molecular Cell 84, 1826–1841.e1825. 10.1016/j.molcel.2024.04.001.

56. Chen, Y., Lyu, R., Rong, B., Zheng, Y., Lin, Z., Dai, R., Zhang, X., Xie, N., Wang, S., Tang, F., et al. (2020). Refined spatial temporal epigenomic profiling reveals intrinsic connection between PRDM9-mediated H3K4me3 and the fate of double-stranded breaks. Cell Research 30, 256–268. 10.1038/s41422-020-0281-1.

57. Tang, S., Hariri, S., Bohn, R., McCarthy, J.E., Koo, J., Pourhosseinzadeh, M., Nguyen, E., Liu, N., Ma, C., Lu, H., et al. (2025). Protecting double Holliday junctions ensures crossing over during meiosis. Nature 647, 776–785. 10.1038/s41586-025-09555-1.

58. Henggeler, A., Orlić, L., Velikov, D., and Matos, J. (2025). Holliday junction–ZMM protein feedback enables meiotic crossover assurance. Nature 647, 766–775. 10.1038/s41586-025-09559-x.

59. Ma, L., O’Connell, J.R., VanRaden, P.M., Shen, B., Padhi, A., Sun, C., Bickhart, D.M., Cole, J.B., Null, D.J., Liu, G.E., et al. (2015). Cattle Sex-Specific Recombination and Genetic Control from a Large Pedigree Analysis. PLoS Genet 11, e1005387. 10.1371/journal.pgen.1005387.

60. Caniçais, C., Sobral, D., Vasconcelos, S., Cunha, M., Pinto, A., Mesquita Guimarães, J., Santos, F., Barros, A., Dória, S., and Marques, C.J. (2025). Transcriptomic analysis and epigenetic regulators in human oocytes at different stages of oocyte meiotic maturation. Developmental Biology 519, 55–64. 10.1016/j.ydbio.2024.12.004.

61. Garcia-Alonso, L., Lorenzi, V., Mazzeo, C.I., Alves-Lopes, J.P., Roberts, K., Sancho-Serra, C., Engelbert, J., Marečková, M., Gruhn, W.H., Botting, R.A., et al. (2022). Single-cell roadmap of human gonadal development. Nature 607, 540–547. 10.1038/s41586-022-04918-4.

62. Powell, T.J., Brown, G.G.B., Allison, R.M., Harper, J.A., Neale, M.J., and Gittens, W.H. (2026). Transcription directs Holliday junction branch migration. bioRxiv, 2026.2003.2013.711646. 10.64898/2026.03.13.711646.

63. Aymard, F., Bugler, B., Schmidt, C.K., Guillou, E., Caron, P., Briois, S., Iacovoni, J.S., Daburon, V., Miller, K.M., Jackson, S.P., and Legube, G. (2014). Transcriptionally active chromatin recruits homologous recombination at DNA double-strand breaks. Nat Struct Mol Biol 21, 366–374. 10.1038/nsmb.2796.

64. Pfister, S.X., Ahrabi, S., Zalmas, L.P., Sarkar, S., Aymard, F., Bachrati, C.Z., Helleday, T., Legube, G., La Thangue, N.B., Porter, A.C., and Humphrey, T.C. (2014). SETD2-dependent histone H3K36 trimethylation is required for homologous recombination repair and genome stability. Cell Rep 7, 2006–2018. 10.1016/j.celrep.2014.05.026.

65. Clouaire, T., and Legube, G. (2019). A Snapshot on the *Cis* Chromatin Response to DNA Double-Strand Breaks. Trends in Genetics 35, 330–345. 10.1016/j.tig.2019.02.003.

66. Baker, Z., Przeworski, M., and Sella, G. (2023). Down the Penrose stairs, or how selection for fewer recombination hotspots maintains their existence. Elife 12. 10.7554/eLife.83769.

67. Singhal, S., Leffler, E.M., Sannareddy, K., Turner, I., Venn, O., Hooper, D.M., Strand, A.I., Li, Q., Raney, B., Balakrishnan, C.N., et al. (2015). Stable recombination hotspots in birds. Science 350, 928–932. 10.1126/science.aad0843.

68. Raynaud, M., Sanna, P., Joseph, J., Clément, J., Imai, Y., Lareyre, J.J., Laurent, A., Galtier, N., Baudat, F., Duret, L., et al. (2025). PRDM9 drives the location and rapid evolution of recombination hotspots in salmonid fish. PLoS Biol 23, e3002950. 10.1371/journal.pbio.3002950.

69. Auton, A., Rui Li, Y., Kidd, J., Oliveira, K., Nadel, J., Holloway, J.K., Hayward, J.J., Cohen, P.E., Greally, J.M., Wang, J., et al. (2013). Genetic Recombination Is Targeted towards Gene Promoter Regions in Dogs. PLOS Genetics 9, e1003984. 10.1371/journal.pgen.1003984.

70. Ramírez, F., Ryan, D.P., Grüning, B., Bhardwaj, V., Kilpert, F., Richter, A.S., Heyne, S., Dündar, F., and Manke, T. (2016). deepTools2: a next generation web server for deep-sequencing data analysis. Nucleic Acids Res 44, W160–165. 10.1093/nar/gkw257.

71. Doran, A.G., Wong, K., Flint, J., Adams, D.J., Hunter, K.W., and Keane, T.M. (2016). Deep genome sequencing and variation analysis of 13 inbred mouse strains defines candidate phenotypic alleles, private variation and homozygous truncating mutations. Genome Biology 17, 167. 10.1186/s13059-016-1024-y.

72. Keane, T.M., Goodstadt, L., Danecek, P., White, M.A., Wong, K., Yalcin, B., Heger, A., Agam, A., Slater, G., Goodson, M., et al. (2011). Mouse genomic variation and its effect on phenotypes and gene regulation. Nature 477, 289–294. 10.1038/nature10413.

73. Lilue, J., Doran, A.G., Fiddes, I.T., Abrudan, M., Armstrong, J., Bennett, R., Chow, W., Collins, J., Collins, S., Czechanski, A., et al. (2018). Sixteen diverse laboratory mouse reference genomes define strain-specific haplotypes and novel functional loci. Nature Genetics 50, 1574–1583. 10.1038/s41588-018-0223-8.

74. Lam, K.-W.G., Brick, K., Cheng, G., Pratto, F., and Camerini-Otero, R.D. (2019). Cell-type-specific genomics reveals histone modification dynamics in mammalian meiosis. Nature Communications 10, 3821. 10.1038/s41467-019-11820-7.

75. Miller, C., Portlock, T., Nyaga, D.M., and O’Sullivan, J.M. (2024). A review of model evaluation metrics for machine learning in genetics and genomics. Front Bioinform 4, 1457619. 10.3389/fbinf.2024.1457619.

76. Kolberg, L., Raudvere, U., Kuzmin, I., Adler, P., Vilo, J., and Peterson, H. (2023). g:Profiler—interoperable web service for functional enrichment analysis and gene identifier mapping (2023 update). Nucleic Acids Research 51, W207–W212. 10.1093/nar/gkad347.

77. An, J.Y., Lin, K., Zhu, L., Werling, D.M., Dong, S., Brand, H., Wang, H.Z., Zhao, X., Schwartz, G.B., Collins, R.L., et al. (2018). Genome-wide de novo risk score implicates promoter variation in autism spectrum disorder. Science 362. 10.1126/science.aat6576.

78. Goldmann, J.M., Wong, W.S., Pinelli, M., Farrah, T., Bodian, D., Stittrich, A.B., Glusman, G., Vissers, L.E., Hoischen, A., Roach, J.C., et al. (2016). Parent-of-origin-specific signatures of de novo mutations. Nat Genet 48, 935–939. 10.1038/ng.3597.

79. Sasani, T.A., Pedersen, B.S., Gao, Z., Baird, L., Przeworski, M., Jorde, L.B., and Quinlan, A.R. (2019). Large, three-generation human families reveal post-zygotic mosaicism and variability in germline mutation accumulation. Elife 8. 10.7554/eLife.46922.

80. Chen, S., Francioli, L.C., Goodrich, J.K., Collins, R.L., Kanai, M., Wang, Q., Alföldi, J., Watts, N.A., Vittal, C., Gauthier, L.D., et al. (2024). A genomic mutational constraint map using variation in 76,156 human genomes. Nature 625, 92–100. 10.1038/s41586-023-06045-0.

81. Agarwal, I., Fuller, Z.L., Myers, S.R., and Przeworski, M. (2023). Relating pathogenic loss-of-function mutations in humans to their evolutionary fitness costs. eLife 12, e83172. 10.7554/eLife.83172.

82. Brick, K., Smagulova, F., Khil, P., Camerini-Otero, R.D., and Petukhova, G.V. (2012). Genetic recombination is directed away from functional genomic elements in mice. Nature 485, 642–645. 10.1038/nature11089.

83. Jiang, Y., Huang, F., Chen, L., Gu, J.H., Wu, Y.W., Jia, M.Y., Lin, Z., Zhou, Y., Li, Y.C., Yu, C., et al. (2023). Genome-wide map of R-loops reveals its interplay with transcription and genome integrity during germ cell meiosis. J Adv Res 51, 45–57. 10.1016/j.jare.2022.10.016.

84. Vergara-Lope, A., Jabalameli, M.R., Horscroft, C., Ennis, S., Collins, A., and Pengelly, R.J. (2019). Linkage disequilibrium maps for European and African populations constructed from whole genome sequence data. Sci Data 6, 208. 10.1038/s41597-019-0227-y.

85. Fang, K., Li, Q., Wei, Y., Zhou, C., Guo, W., Shen, J., Wu, R., Ying, W., Yu, L., Zi, J., et al. (2021). Prediction and Validation of Mouse Meiosis-Essential Genes Based on Spermatogenesis Proteome Dynamics. Mol Cell Proteomics 20, 100014. 10.1074/mcp.RA120.002081.

86. Guo, S., Zhang, Y., Fei, C., Liu, X., Xia, W., Luo, M., Wei, G., Qin, W., Xiong, C., Li, H., et al. (2025). Deciphering meiotic chromatin organization by SYCP3. Nucleic Acids Research 53. 10.1093/nar/gkaf460.

87. Wang, R.J., Dumont, B.L., Jing, P., and Payseur, B.A. (2019). A first genetic portrait of synaptonemal complex variation. PLoS Genet 15, e1008337. 10.1371/journal.pgen.1008337.

88. Reynolds, A., Qiao, H., Yang, Y., Chen, J.K., Jackson, N., Biswas, K., Holloway, J.K., Baudat, F., de Massy, B., Wang, J., et al. (2013). RNF212 is a dosage-sensitive regulator of crossing-over during mammalian meiosis. Nat Genet 45, 269–278. 10.1038/ng.2541.

